# A Total-Group Phylogenetic Metatree for Cetacea and the Importance of Fossil Data in Diversification Analyses

**DOI:** 10.1101/2020.06.24.169078

**Authors:** Graeme T. Lloyd, Graham J. Slater

## Abstract

Phylogenetic trees provide a powerful framework for testing macroevolutionary hypotheses, but it is becoming increasingly apparent that inferences derived from extant species alone can be highly misleading. Trees incorporating living and extinct taxa are are needed to address fundamental questions about the origins of diversity and disparity but it has proved challenging to generate robust, species–rich phylogenies that include large numbers of fossil taxa. As a result, most studies of diversification dynamics continue to rely on molecular phylogenies. Here, we extend and apply a recently developed meta–analytic approach for synthesizing previously published phylogenetic studies to infer a well–resolved set of species level, time–scaled phylogenetic hypotheses for extinct and extant cetaceans (whales, dolphins and allies). Our trees extend sampling from the ∼ 90 extant species to over 400 living and extinct species, and therefore allow for more robust inference of macroevolutionary dynamics. While the diversification scenarios we recover are broadly concordant with those inferred from molecular phylogenies they differ in critical ways, most notably in the relative contributions of extinction and speciation rate shifts in driving rapid radiations. Supertrees are often viewed as poor substitute for phylogenies inferred directly from character data but the metatree pipeline overcomes many of the past criticisms leveled at these approaches. Meta–analytic phylogenies provide the most immediate route for integrating fossils into macroevolutionary analyses, the results of which range from untrustworthy to nonsensical without them.

---

It is now widely accepted that a phylogenetic framework is essential for addressing questions regarding diversification dynamics, phenotypic evolution, and historical biogeography. The covariances between species that are imposed by the hierarchical structure of a phylogenetic tree mean that any attempt to understand the processes responsible for generating observed patterns of diversity must take the tree and its associated branch lengths into account (Felsenstein, 1985; Harvey and Pagel, 1991; Foote, 1996; O’Meara et al., 2006; Ree and Smith, 2008). As a consequence of this phylogenetic dependence, the development of new tools for inferring macroevolutionary dynamics has been paralleled by innovations in the field of phylogenetic inference, and it is now possible to infer time-scaled trees using complex models of molecular evolution applied to genome-scale data.

The need for a well–resolved, time–calibrated phylogeny places substantial constraints on the kinds of clades that are accessible to most biologists for testing macroevolutionary hypotheses. Some authors have noted that clades are often selected for study due to their tractability rather than because they are suitable candidates for testing a particular hypothesis, resulting in a form of empirical ascertainment bias (Beaulieu and O’Meara, 2018, 2019). For example, early burst models of adaptive radiation arose to explain the origins of higher taxa (Simpson, 1944, 1953; Van Valen, 1971; Valentine, 1980; Humphreys and Barraclough, 2014; Slater and Friscia, 2019) but have mostly been tested in lower level clades, such as genera, where the early burst signal is conspicuously lacking (e.g., Harmon et al., 2010). Although lower level clades certainly have a role to play in comparative biology (Schluter, 2000; Losos, 2009; Donoghue and Edwards, 2019), there is a pressing need to develop suitable phylogenetic frameworks for studying macroevolutionary pattern and process at higher taxonomic levels.

The major barrier to obtaining appropriate phylogenetic frameworks for higher–level clades has always been data availability (Smith et al., 2009). The “Supermatrix” approach was initially suggested as a solution to this problem (Sanderson et al., 1998; Gatesy et al., 2002; de Queiroz and Gatesy, 2007). Here, one obtains all available sequence data for a clade of interest through a combination of direct sequencing and from repositories such as Genbank. Sequences are aligned and concatenated to create a large but sparsely sampled matrix that can be analyzed using standard phylogenetic software and methods. Concerns regarding the impact of missing data and data quality (e.g., McMahon and Sanderson, 2006) have, more recently, led to alternative approaches based on bioinformatic pipelines Smith et al. (2009) or patching of subclades onto backbone trees (Jetz et al., 2012; Tonini et al., 2016; Jetz and Pyron, 2018; Upham et al., 2019). These methods have proved effective for generating large, higher–level phylogenetic hypotheses (particularly where taxonomic information can also be used to constrain the placement of species that lack character data) and have yielded novel insights into diversification dynamics, trait evolution and historical biogeographic patterns. Recent examples include a 5,284 species tree of agariomycete fungi (Varga et al., 2019), an 11,638 species tree of extant fishes (Rabosky et al., 2018), and a 353,185 species tree of seed plants (Smith and Brown, 2018).

While these methods provide promise for extant clades, they cannot be used to generate phylogenetic hypotheses for most of the > 99% of life that is now extinct (Raup, 1994). This is particularly problematic given that fossil data play a critical role in refining estimates of ancestral character states (Finarelli and Flynn, 2006), choosing among competing models of trait evolution (Slater et al., 2012), inferring ancestral biogeographic patterns (Meseguer et al., 2014), and understanding speciation and extinction dynamics through time (Mitchell et al., 2018; Louca and Pennell, 2020). The difficulty in generating large character–taxon matrices for fossil taxa is due in large part to the unique and often subjective ways in which morphological characters and their states are defined and coded across studies. Unlike molecular data, where character states are universally coded, two morphological matrices with partially overlapping taxon lists cannot be concatenated without extensive revision of characters and re–coding of their states, which is, in itself, a challenging, time–consuming, and potentially impossible task. The effect of this incompatibility is that, although the number of species included in morphological character–taxon matrices has continued to increase over the past few decades (fig 1), they lag well behind molecular datasets in size. One recent study included character state codings for 501 OTUs (Hartman et al., 2019), but this is twice the size of the next largest matrix published to date (N=254, Mo et al., 2012).

**Fig. 1.**
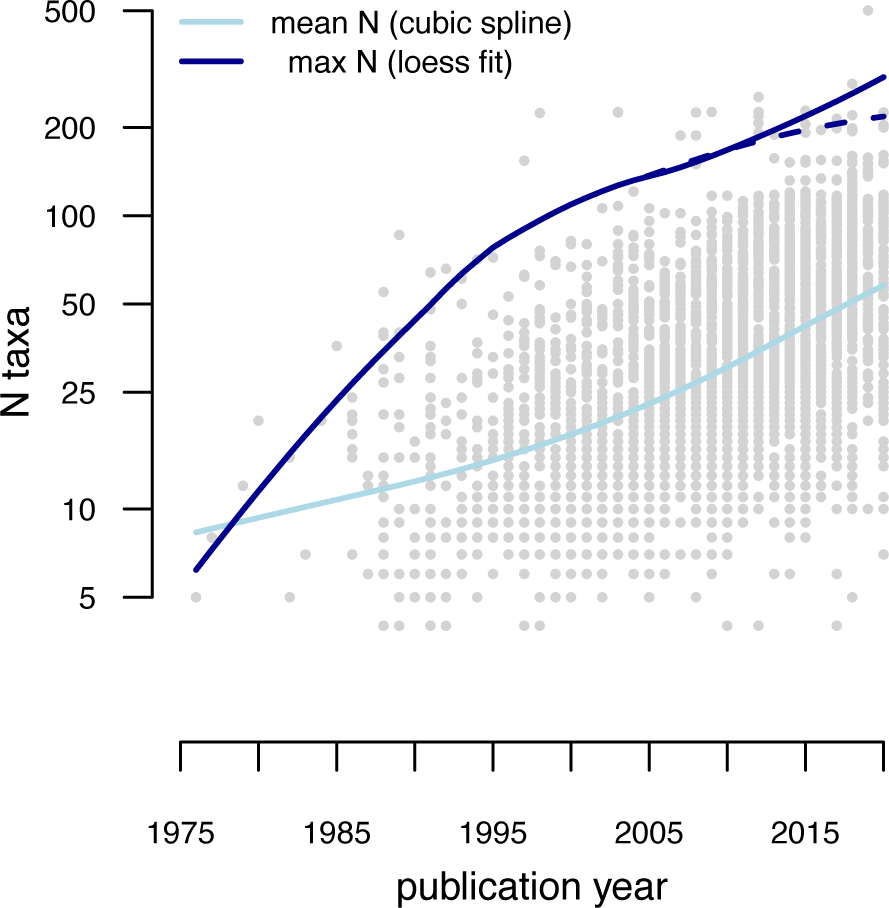
Although the number of taxa included in morphological character–taxon matrices has increased over time, they lag behind the largest molecular datasets. Based on a cubic spline (light blue line) fitted to log(number of taxa) in 3671 morphological studies (graemetlloyd.com/matr.heml), the average dataset has only increased from 8.3 OTUs in 1975 to 58 OTUs in 2020 (note log scale on y–axis). The maximum number of taxa has also increased, corroborated by a loess fit (dark blue solid line). Removing Hartman et al. (2019), which contains the largest number of taxa by a factor of 2, indicates a slow–down in the rate of increase towards the present (dark blue dashed line).

Supertree methods provide an alternative avenue for the inference of large phylogenies of extinct taxa. Supertrees are a class of consensus tree in which a set of topologies derived from distinct datasets are summarized in some common form to yield a topology containing shared or well–supported splits (Sanderson et al., 1998; Sanderson and Driskell, 2003; Bininda-Emonds et al., 2002; Bininda-Emonds, 2004). Importantly, supertree methods can accommodate sets of input trees with partially or non–overlapping leaf sets, and they therefore provide a way of synthesizing morphological character–taxon matrices covering distinct clades without re–coding characters or concatenating matrices. The best–known method for combining trees is Matrix Representation with Parsimony (MRP), where all input topologies are represented using a binary coding scheme (Fig. 2). Each column, or character, in a MRP matrix represents a bipartition from one of the source trees. An entry of “1” for a given row indicates the presence of that taxon within the clade, “0” indicates its exclusion from the clade, and “?” indicates that the taxon is not represented in the source tree in question (Baum, 1992; Ragan, 1992; Baum and Ragan, 2004, Fig 2C). A supertree containing the union of tips over the source trees may then be inferred using standard parsimony methods. Like supermatrices, supertrees (and MRP supertrees in particular) have been criticized on a number of grounds. Character non–independence necessarily arises due to reuse of characters across multiple analyses (Springer and de Jong, 2001), and issues concerning the relative quality of individual studies must also be addressed (Gatesy et al., 2004). Further sources of concern include how to select and code topologies produced from analysis of the same matrix (Gatesy et al., 2004), weighting of strongly versus weakly supported nodes (Gatesy and Springer, 2004), the potential recovery of clades that are not found in any of the input trees (Pisani and Wilkinson, 2002; Bininda-Emonds, 2003; Wilkinson et al., 2005) and how best to deal with supraspecific OTUs (Page, 2004).

**Fig. 2.**
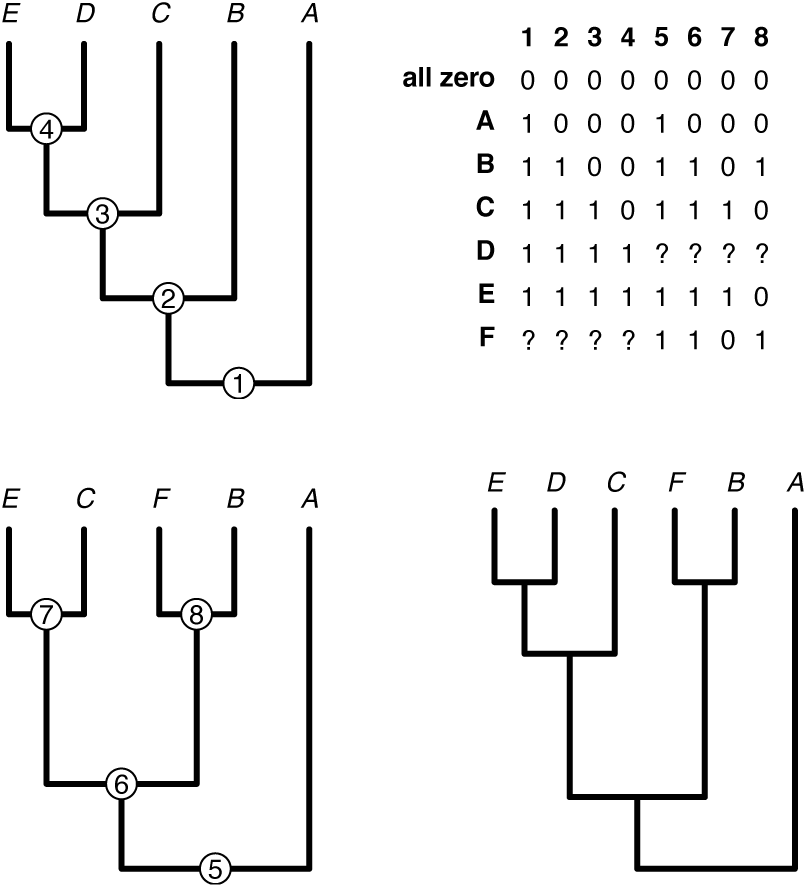
Matrix representation of tree topologies allows for the inference of total clade trees even if taxa are missing from individual source trees. Here, the two source trees on the left differ in that tree 1 does not sample taxon F while tree 2 is missing taxon D. Parsimony analysis of the matrix representation of the two topologies results in a single shortest tree that captures the intuitive relationships of F with B and C as sister to D+E that are not both present in either of the two source trees.

In response to these criticisms, a number of alternative supertree approaches have been developed (e.g, Bininda-Emonds, 2004; Semple et al., 2004; Levasseur and Lapointe, 2006; Steel and Rodrigo, 2008; Lin et al., 2009; Ranwez et al., 2010; Swenson et al., 2012; Akanni et al., 2015; Kettleborough et al., 2015; Fleischauer and Böcker, 2017). Some of these methods allow for character weighting based on information such as bootstrap values, or relative importance of a given source tree, but none provides a straightforward way to explicitly accommodate phylogenetic uncertainty within individual source studies, particularly where the number of studies is large. Furthermore, none of these approaches or pipelines explicitly deal with earlier criticisms of the supertree paradigm that are rooted in issues of data reuse and redundancy (Springer and de Jong, 2001; Gatesy and Springer, 2004; Gatesy et al., 2004). Lloyd et al. (2016) introduced an alternative approach that they called a “metatree”. As with MRP supertrees, metatrees use binary encoding of tree topologies to generate a matrix that can be analyzed using standard phylogenetic methods. The principal difference between an MRP supertree and a metatree is that the matrix from which MRP supertrees are built typically uses individual tree topologies gleaned from published papers as data, while the metatree approach explicitly requires reanalysis of morphological character-taxon matrices to sample and encode *all* optimal topologies. Moreover, the metatree pipeline introduces specific rules to ameliorate concerns associated with data redundancy and uncertainty in the inference of source trees. In practice, metatrees tend to lead to more resolved consensus topologies than traditional MRP supertrees (compare Lloyd et al. 2016 to Lloyd et al. 2008) while also better accommodating phylogenetic uncertainty in the source studies than the figured trees typically used by supertree methods (Bell and Lloyd, 2015).

In this paper we leverage the metatree approach to assess diversification dynamics in extant and extinct cetaceans (whales, dolphins and relatives). A number of recent studies based on molecular phylogenies have provided evidence for a recent increase in cetacean net diversification rates during the past 10 Ma, driven by rapid speciation of ocean dolphins (Delphinidae) (Steeman et al., 2009; Slater et al., 2010; Rabosky, 2014; Rabosky and Goldberg, 2015). However, the relative contributions of speciation and extinction rate variation to trends in net diversification can be extremely difficult to disentangle using phylogenies of extant taxa (Liow et al., 2010; Louca and Pennell, 2020) and the rich cetacean fossil record suggests that different dynamics may have been at play during the past 36 million years than might be suggested on the basis of molecular phylogeny alone (Quental and Marshall, 2010; Morlon et al., 2011; Marx and Fordyce, 2015). Until now, the lack of a densely–sampled higher–level phylogeny of the clade has precluded thorough comparison of diversification dynamics inferred from molecular and fossil phylogenies. We here use the metatree pipeline (Lloyd et al., 2016) to assemble a comprehensive set of phylogenetic hypotheses for extant and extinct cetaceans. We then use a Bayesian model–averaging approach (fossilBAMM Rabosky et al., 2014; Mitchell et al., 2018) to estimate rates of speciation and extinction through time and across branches of the time–scaled cetacean trees. Our results demonstrate that simultaneous analysis of extinct and extant taxa can yield different conclusions regarding macroevolutionary dynamics than are derived from analyses of extant taxa alone, and stress the important of paleo–phylogenetic approaches for studying macroevolutionary dynamics.

## Materials and Methods

The metatree approach was fully described in Lloyd et al. (2016) but we provide an overview here in the context of assembly of our cetacean metatree. For ease of reference, our pipeline is summarized visually as a flow–chart in Figure 3 and our description of methods follow this structure.

**Fig. 3.**
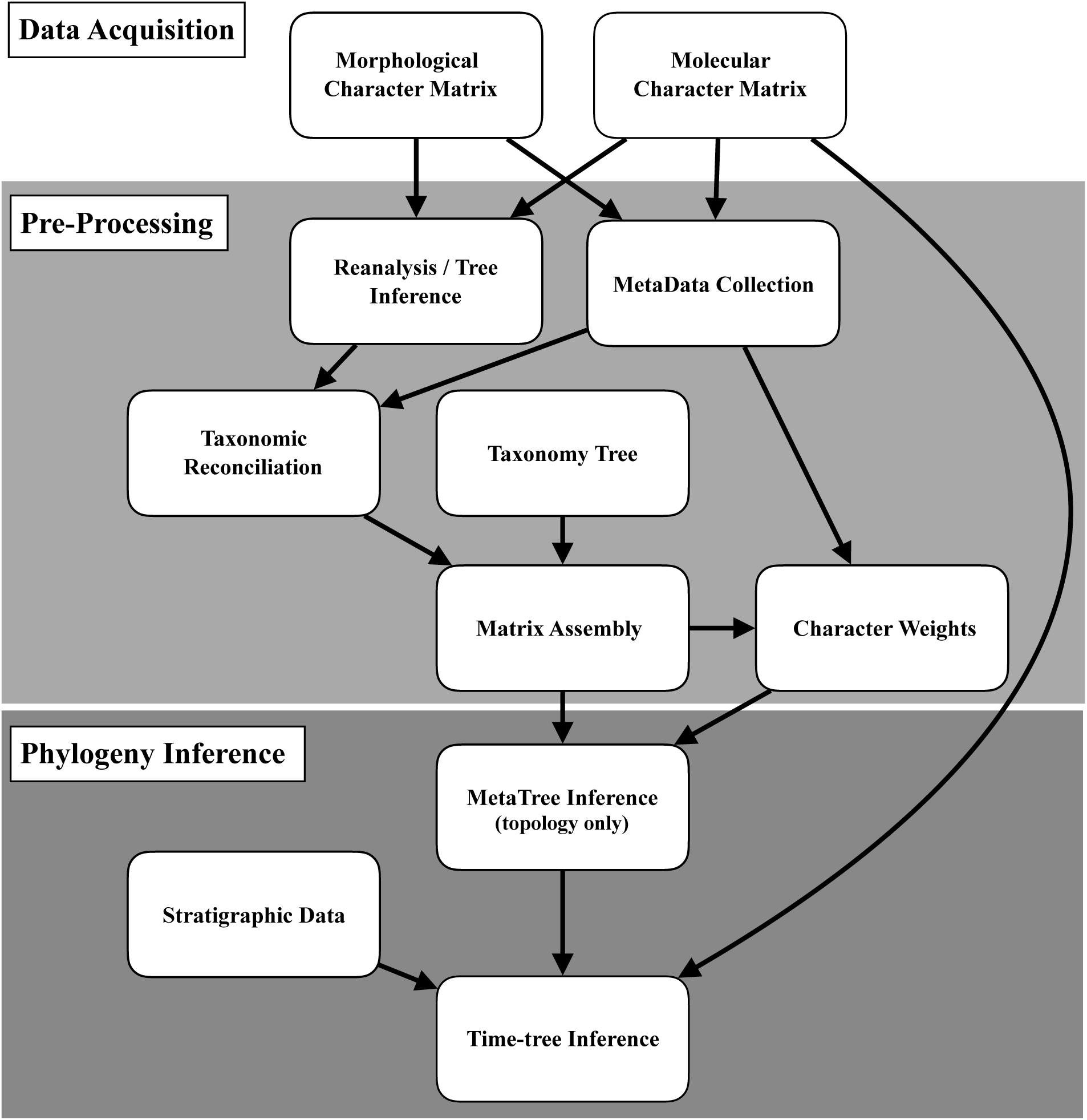
Schematic showing the general outline of metatree assembly. Description of methods used in the assembly of our cetacean metatrees follow this workflow.

### Data Acquisition

#### Morphological character data

We collected 146 morphological character matrices from 143 published studies (See Supplementary Bibliography). Sampled studies range in publication date from 1994 to 2020. New species of (typically extinct) cetacean are described with sufficient regularity that such a tree can quickly become out-of-date. However, our pipeline allows easy integration of additional data for continuous updating. We included phylogenetic analyses of exclusively extinct, exclusively extant, and both extinct and extant taxa in our dataset. The only requirement for inclusion was that a morphological character matrix was provided in the paper, the associated supplementary methods, or on some repository such as Morphobank (O’Leary and Kaufman, 2011). All character matrices have been deposited on the cetacean metatree GitHub repository (https://github.com/graemetlloyd/ProjectBlackFish). We retained information regarding any character weighting or ordering schemes used in the published analyses. To minimize the impact of data duplication, we removed molecular data, where included, from each alignment.

#### Molecular character data

Molecular evidence may provide a strong and divergent phylogenetic signal for extant taxa compared with the signal provided by morphological data. Parsimony analyses of morphological character data often employ a molecular scaffold approach, whereby tree searches are constrained to recover topologies for extant taxa that are consistent with molecular estimates. However, the fact that a single topology constraint must typically be enforced means topological uncertainty inherent in tree inference from molecular data cannot be accommodated. The metatree pipeline can readily incorporate molecular data as an additional data source, thus accommodating topological uncertainty. We here used the molecular supermatrix of McGowen et al. (2009). This matrix contains 42,335 characters from 45 nuclear loci, mitochondrial genomes, and transposon insertion events coded for 91 taxa (four artiodactyl outgroups and 87 of 91 currently recognized cetacean species).

### Pre-processing

#### Reanalysis

We reanalyzed each morphological character–taxon matrix, under the outgroup, weighting, and ordering schemes specified by the original authors, using the maximum parsimony software TNT (Goloboff et al., 2008). The use of parsimony allowed us to balance our desire to incorporate phylogenetic uncertainty via a set of most parsimonous trees with the need for an efficient pipeline for processing large numbers of source datasets while guaranteeing convergence on an optimal set of solutions. However, any approach (Maximum Likelihood, Bayesian Inference) could be used to reanalyze source data. The settings for each analysis were determined on the basis of matrix size. For 24 or fewer taxa the implicit enumeration option was used, which guarantees that all optimal topologies will be returned. For 25 or more taxa, 20 separate replicates of TNT’s “New Technology Searches” were performed, each starting with a random seed. Trees from each replicate were then combined and a final round of tree bisection–reconnection was performed. In some cases the maximum tree limit (100,000) was hit, indicating additional equally optimal relationships exist. In order not to miss these topologies, searches were repeated until the least frequent bipartition was found at least twice suggesting complete coverage was reached). For the purposes of inference, trees were summarised using matrix representation, with all duplicate bipartitions removed. All unique bipartitions were equally weighted as under parsimony there is no clear basis for considering bipartition frequency within the set of shortest trees as a measure of support.

The molecular data from McGowen et al. (2009) were analyzed in MrBayes v 3.2 (Ronquist et al., 2012) via the CIPRES portal (Miller et al., 2010) using the same MCMC settings and models as in the original study. In Bayesian phylogenetic methods, topologies (and their associated branch lengths) are sampled in proportion to their posterior probabilities, which means that bipartition frequency is meaningful. We encoded unique bipartitions found within 1000 trees drawn at random from the posterior sample using matrix representation but assigned each column of the resulting character matrix a weight corresponding to its frequency in the posterior sample. For example, a clade sampled in only 10% of the posterior sample was assigned a weight that is one–tenth that of a clade present in all trees in the sample.

#### Metadata

We recorded two key pieces of metadata from each source study. First, we noted whether supra–specific Operational Taxonomic Units (OTUs) in the character matrix were coded from a specific taxon or taxa. For example, the NEXUS file of Godfrey et al. (2017) lists the OTU *Phocageneus* but the paper itself confirms that this OTU is *Phocageneus venustus*. If a supra–specific OTU was coded from more than one taxon, all were recorded if listed in the paper. If no species–level taxa were listed, we retained the supra–specific taxon as the unit of analysis. Direct editing of names in NEXUS files can lead to problems if taxa are later synonymized or names are altered. We therefore generated custom XML files for each source study in which each OTU was reconciled to its constituent taxa, as recorded in the Paleobiology Database (Peters and McClennen, 2016). Cetaceans, both extinct and extant, are extremely well documented in the Paleobiology Database thanks largely to the efforts of Mark Uhen (e.g., Uhen and Pyenson, 2007).

Additionally, as the database is continually updated and has an API (Peters and McClennen, 2016), dynamic updating of taxonomy can be achieved in future metatree iterations. Seventy undescribed OTUs did not have a species name. Following Lloyd et al. (2016) we retain these as “valid” taxa in our analyses because 1) they may represent key data for resolving phylogenetic relationships, dating the tree, or performing downstream macroevolutionary analysis, and 2) there is a clear and repeated tendency for these specimens to become the holotypes of new species (for example, the specimen-level OTU, “GSM 109” was included in multiple trees prior to its formal description as *Echovenator sandersi* by Churchill et al. 2016). Such OTUs are a common feature in other clades too; 10% (97 of 961) of OTUs in Lloyd et al. (2016) were unnamed specimens.

The second piece of information noted was whether the matrix was based on a previous study. For example, the matrix of Godfrey et al. (2017) was based on an earlier study by Lambert et al. (2014) and was itself later used by Boersma et al. (2017). This information was added to each matrix’s XML file for use in determining relative weighting or study redundancy when assembling the final MRP matrix.

#### Taxonomic reconciliation and taxonomy tree

Before matrix representations for each source study can be combined to form a global matrix, a list of valid species must be decided on and taxonomic assignments reconciled to this list. We dynamically reconciled tip names recorded in the XML files to currently valid taxon names recognized in the Paleobiology Database, for example updating junior synonyms to their senior synonyms. This procedure allows taxonomic information to be automatically updated and limits human error while updating names.

The taxonomic hierarchy present in the database also represents a pseudo-phylogenetic hypothesis (Soul and Friedman, 2015), a feature exploited here in two ways. First, supraspecific taxon rows that cannot be reconciled to specific taxa can be replaced with a set of duplicated rows corresponding to species-level OTUs, avoiding the situation where, for example, *Balaenoptera* and *Balaenoptera musculus* exist as separate tips in the final metatree. Second, the taxonomy can be included as an additional, albeit a heavily down–weighted data set (Gatesy and Springer, 2004). This is important as the presence of a basic but comprehensive estimate of phylogeny derived from taxonomy can ameliorate inference issues that might arise due to a lack of data overlap (an affliction of formal supertrees often termed “rogue taxa”). For example, Mysticetes and Odontocetes should logically be separated as clades, but if all phylogenetic analyses only focused on one or the other of these clades, then information on their reciprocal monophyly would be lacking.

The use of a taxonomy tree also allows for the inclusion of species that are un–sampled in the set of source trees. Higher–level analyses of extant clades have included taxa for which molecular data are unavailable by simultaneously enforcing topology constraints, based on taxonomy, and integrating over possible placements of missing taxa under a birth–death process (e.g., Kuhn et al., 2011; Jetz et al., 2012; Rabosky et al., 2018; Upham et al., 2019). We here increased taxonomic coverage for fossil cetaceans by producing two additional versions of our MRP matrix, one in which all species assigned to a sampled genus were included and another where all species assigned to Cetacea in the Paleobiology Database were included. We refer to these analyses as GENUS and ALL, respectively, for the remainder of this paper, with the species–level analysis referred to as EXCLUDE to account for the fact that unsampled OTUs were excluded. It should be stressed that, because our pipeline treats taxonomic bipartitions as data that are down–weighted relative to bipartitions derived from phylogenetic analyses, this approach does not force a taxonomic structure on the result where there are primary character data available that disagree with it (cf. Jetz et al., 2012; Rabosky et al., 2018; Upham et al., 2019).

#### Matrix assembly and character weighting

After taxonomic reconciliation, it is straightforward to merge MRP matrices from source studies into a global character–taxon matrix. At this stage, we also compute character weights, based on three attributes of source studies: non–independence, date of publication, and size (measured in number of MRP characters). Older character matrices that were reused in a subsequent study without modification of the characters themselves (e.g., by the addition of a single new taxon to an existing dataset) were deemed redundant and automatically removed. The remaining non–independent data sets were assigned equal weights that sum to one, with this weight being applied to each character of the data set. Next, following Lloyd et al. (2016), publication year weights were assigned such that the oldest included data–set received a weight of 10 (an order of magnitude higher than the weight assigned to the taxonomy tree) and with weights doubling every two years. Again, this weight was applied to each character in a data set. Finally, some data sets generated more MRP characters than others simply because they contained greater phylogenetic uncertainty and without intervention these would dominate the final tree. To account for this individual characters (bipartitions) were weighted such that any within-data-set characters (biparitions) with which they conflict are clustered and down-weighted such that they sum to one, with any unconflicted characters being weighted one. These different character weights were combined by taking the product of the three criteria–based weights. Other ways of weighting characters and source studies are possible (interested readers can consult the *in development metatree* R package github.com/graemetlloyd/metatree for more information and options) but we have so far found the above to work well across a range of groups.

TNT (Goloboff et al., 2008) requires that weights fall in the range 0.50 – 1000.00. We set default weights of one for the taxonomy tree (always enforced) and 10 for the minimum phylogenetic weight, but no initial maximum weight can be specified and in practice this may exceed 1000.00. When this occurred, we rescaled the phylogenetic weights only to fall on a 10.00 – 1000.00 scale. Lloyd et al. (2016) applied multistate characters to effectively stretch this maximum possible weight to 31000.00, but we found that this dramatically slowed the run time of TNT. Thus when it came time to upweight molecular topologies relative to morphological topologies, we instead assigned them maximum weights combined with column (character) duplication. Column duplication is identical to numerically upweighting character state changes or site likelihoods using integer–valued weights, and is the method employed in model–based phylogenetic inference tools such as RAxML (Stamatakis, 2014) and BEAST 2 (Bouckaert et al., 2014).

### Phylogeny Inference

#### Metatree inference

Prior to analysis, the final MRP matrix was subjected to Safe Taxonomic Reduction (STR: Wilkinson, 1995) using the `SafeTaxonomicReduction`() function in the R (R Development Core Team, 2019) package `Claddis` (Lloyd, 2016) and an all–zero outgroup was added to provide character polarity during the tree search (Baum, 1992; Ragan, 1992). We performed 1000 independent parallel tree searches using TNT with the `xmult` option for multiple replications using sectorial searches, drifting, ratchet and fusing invoked at level 10, and a maximum of 1000 trees held in memory (Goloboff et al., 2008). We reinserted STR taxa using the `SafeTaxonomicReinsertion` function in `Claddis` (Lloyd, 2016) and constructed a strict consensus tree from the final sample of shortest trees.

#### Time-tree Inference

The result of metatree inference is a set of most parsimonious topologies that can be summarized using consensus methods. However, macroevolutionary analyses require topologies with associated branch lengths in units of time. Paleontological approaches for time scaling phylogenies have historically been somewhat arbitrary in nature (for a review see Hunt and Slater, 2016). However, these approaches have recently been superseded by probabilistic methods that allow for simultaneous inference of topology and branch lengths for extinct and extant species under a birth–death process (Heath et al., 2014; Gavryushkina et al., 2014, 2017).

We combined three sources of data to sample a distribution of time–scaled phylogenies for extinct and extant cetaceans using BEAST 2.5.2 (Bouckaert et al., 2014). We first used the strict consensus metatree topology to derive a series of topological constraints for each BEAST analysis. No character data were used for extinct taxa and so no morphological clock was invoked to derive branch lengths. In an analysis of extant taxa only, the resulting topological arrangements among unconstrained taxa would be random, but for extinct taxa they are influenced by stratigraphic age, via the use of the Fossilized birth–death process tree prior (Heath et al., 2014; Gavryushkina et al., 2014, 2017). As a result, sampled topologies can be thought of as reflecting a balance between strong prior belief, in the form of hard topological constraints derived from metatree inference, and stratigraphic data. For each extinct terminal taxon in our strict consensus topology, we first queried the Paleobiology Database to obtain the age of first occurrence. The age of each taxon was then specified as the beginning and end dates for the stage of first occurrence, based on the 2018 International Commission on Stratigraphy updated chronostratigraphic chart (http://stratigraphy.org/ICSchart/ChronostratChart2018-07.pdf). Where possible, we supplanted PBDB–derived ages, with more refined biostratigraphic or radiometric age estimates taken from primary sources or previous phylogenetic analyses and revisions (Table S1).

We selected a subset of the alignment from McGowen et al. (2009) for use in our BEAST analyses. Molecular data can provide important information regarding the relative branch lengths for extant taxa, particularly in clades lacking fossil representatives. Preliminary attempts to perform BEAST analyses using the entire alignment yielded poor mixing, even after very long (> 10^8^ generations) runs. We therefore used SortaDate (Smith et al., 2018) to identify and rank genes that were most congruent with the topology reported by McGowen et al. (2009) and that displayed the most clock–like behavior. Based on these criteria *Cytochrome B* was identified as the most appropriate gene and was used in Bayesian estimation of topology and branch lengths (note that the same gene was used, due to its availability for all 87 extant taxa, in McGowen et al., 2009) We determined that an uncorrelated relaxed clock with log–normally distributed rates best fitted the molecular data, based on comparison of marginal likelihoods computed for a fixed topology of extant taxa (see Supplementary Information). We set informative priors on the net diversification (*r* ∼ exponential[1.0]), and relative extinction rates (*ϵ* ∼ *β*[2.0, 1.0] based on Marshall’s 2017 third paleobiological law that speciation ≈ extinction), and placed a flat prior on fossilization probability (*s* ∼ U[0, 1]). For the origin of the FBD process we specified an offset exponential prior, with an offset of 54 million years, corresponding to the age of the oldest known cetacean *Himalayacetus subanthuensis* (Bajpai and Gingerich, 1998), and mean of 3.5 that resulted in a 95% quantile corresponding to the Cretaceous – Paleogene boundary (66 Ma). We ran two chains for 10^8^ generations, sampling every 10^5^ generations and, after visually checking for convergence and parameter effective sample sizes > 200 using Tracer v1.7.1, we discarded a chain–specific burn–in and combined tree files. Attempts to produce a maximum clade credibility tree annotated with mean or median branch lengths failed due to negative branch lengths, indicating conflict between the most frequently sampled topology and the distribution of underlying branch lengths. Instead, we sampled the Maximum *A Posteriori* (MAP) tree for visualization purposes and for subsequent macroevolutionary analysis.

### Inference of Diversification Dynamics

Analyses based on molecular phylogenies of extant cetacean phylogeny have recovered evidence for an increase in mean net diversification rates during the past 10 Ma, driven by increased rates of speciation in oceanic dolphins (McGowen et al., 2009; Steeman et al., 2009; Slater et al., 2010; Rabosky et al., 2014). It is well known that inference of trait evolution dynamics can be misleading when based on phylogenies of extant taxa alone (Finarelli and Flynn, 2006; Slater et al., 2012, 2017) and some evidence suggests that inference of cetacean diversification dynamics may suffer from similar issues (Quental and Marshall, 2010; Morlon et al., 2011). We used fossilBAMM (Mitchell et al., 2018) to infer speciation and extinction dynamics for each of the MAP time–scaled cetacean metatrees. As with the standard form of BAMM (Rabosky et al., 2014), fossilBAMM is a Bayesian model–averaging approach that samples speciation and extinction rates along branches of a phylogentic tree while allowing for shifts in one or both rates. The method requires a bifurcating, time–scaled tree containing living and extinct taxa, as well as the number of unique fossil occurrences for tips included in the tree. We queried the Paleobiology Database to recover unique stratigraphic occurrences associated with each terminal taxon present in each of the three metatrees. Taxa not in the database (i.e., undescribed taxa) were treated as having unique single occurrences. Prior to analysis, we also pruned sampled ancestors from the MAP trees to avoid biasing estimates of speciation and extinction due to very short terminal edges. We determined priors for each analysis by using the `setBAMMpriors` function in the `BAMMtools` library (Rabosky et al., 2014) and ran two independent MCMC chains for 10^8^ generations, sampling every 10^4^. We checked for convergence and large effective sample sizes using functions in the `coda` library (Plummer et al., 2006) and processed post-burnin output using functions from the `BAMMtools` library (Rabosky et al., 2014). To compare and contrast trends in diversification dynamics derived from the three metatrees, we plotted median and 95% confidence intervals for speciation, extinction and net diversification rates through time using the `plotRateThroughTime`() function. To compare branch and clade specific rates, we also plotted mean per–branch rates on the respective phylogenetic hypotheses.

## Results

### Metatree Inference

Analysis of the 147 source studies (146 morphological plus 1 molecular) resulted in an MRP matrix comprising 494 species, approximately two-thirds of all recognized cetacean taxa, and 14257 binary characters. Safe Taxonomic Reduction reduced the size of the matrix for analysis to 440 taxa. The strict consensus of 1000 most parsimonious trees, after reinsertion of STR tips, is remarkably well–resolved (78% of nodes, Fig 4a) with polytomies concentrated in basilosaurid archaeocetes, Balaenopteroidea (including extant rorquals), squalodontid odontocetes, and the beaked whale genus *Mesoplodon*. Adding taxa to the taxonomy source tree allowed us to increase taxonomic coverage but, without additional data to place the new species, tended to lead to much less well resolved strict consensus topologies. Specifically, the percentage of resolved nodes dropped to 56% in the GENUS tree (615 taxa, 14326 characters; Fig 4b) and to 29% for the ALL tree (746 taxa, 14344 characters; Fig 4c).

**Fig. 4.**
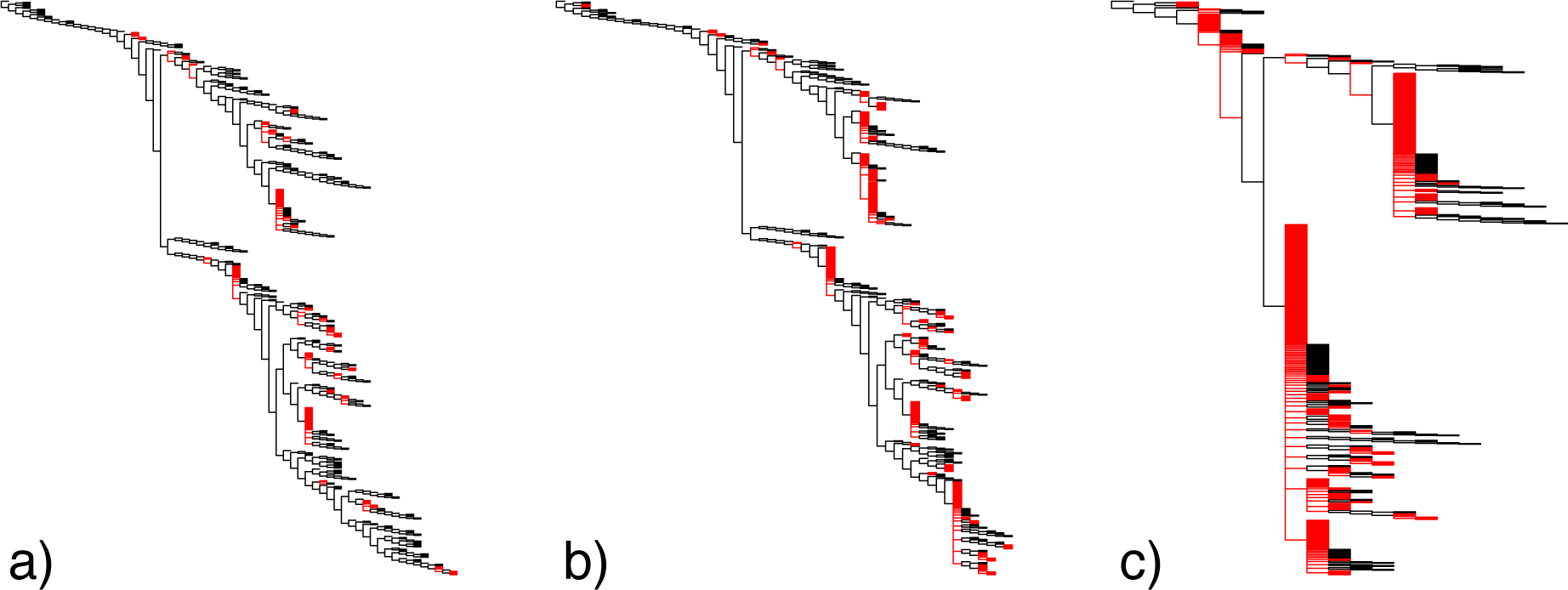
Strict Consensus metatrees of cetaceans based on inclusion of a) 494 species–level OTUs from source trees, b) 615 species belonging to genus–level OTUs from source trees, and c) all 746 valid cetacean taxa recorded in the Paleobiology Database. Black edges represent those branches arising from bifurcating nodes. Red edges are those arising from polytomies either due to uncertainty in the MPT set or due to reinsertion of taxa that were removed prior to analysis during safe taxonomic reduction.

The lack of resolution in the ALL strict consensus metatree, in particular, poses problems for reliable inference of topology and branch lengths during time–tree inference. We therefore used Matrix Representation with Likelihood (MRL; Nguyen et al., 2012) to generate a more stable estimate of topology for this dataset. We first switched the codings of 50% of data columns selected at random, such that “0” became “1” and vice versa to avoid violating the assumptions of the symmetric Markov models employed in phylogenetics software. We then used RAxML v. 8 (Stamatakis, 2014) to find the maximum likelihood estimate of topology under the BINCAT model with rate heterogeneity disabled (−V option). Taxa removed during safe taxonomic reduction were subsequently reinserted and the resulting tree was then used as a topology constraint for BEAST analyses.

### Time-tree Inference

The availability of stage–level or finer stratigraphic data reduced the number of taxa from 494 to 487 for Bayesian estimation of topology and branch lengths for the EXCLUDE dataset. Divergence time estimates among extant clades in the MAP tree (Fig 5) are broadly consistent with previously published estimates for cetaceans but the inclusion of extinct taxa yields novel insights into the timescale of whale evolution. The cetacean stem extends back to 51.3 Ma, with the divergence of the semi–aquatic archaeocete clade Pakicetidae (*Pakicetus, Ichthyolestes*, and *Nalacetus*) from all other cetaceans. Fully aquatic cetaceans (the Pelagiceti of Uhen, 2008) originate at 41.7 Ma with the divergence of a clade comprising the paraphyletic Basilosauridae and Neoceti. The time–scaled metatree emphasizes that many of the gaps along long internal branches of molecular time–trees should be filled with now–extinct radiations. For example, the long stem lineage leading to crown odontocetes that is implied by molecular phylogenies is filled in by the radiations of Xenorophidae, Waipatiidae, Patricetidae, and Squalodontidae. The former diversity of Physeteroidea and Platanistoidea is also apparent in the time–scaled tree, despite the low diversity of these clades in modern times (3 and 1 extant species, respectively).

**Fig. 5.**
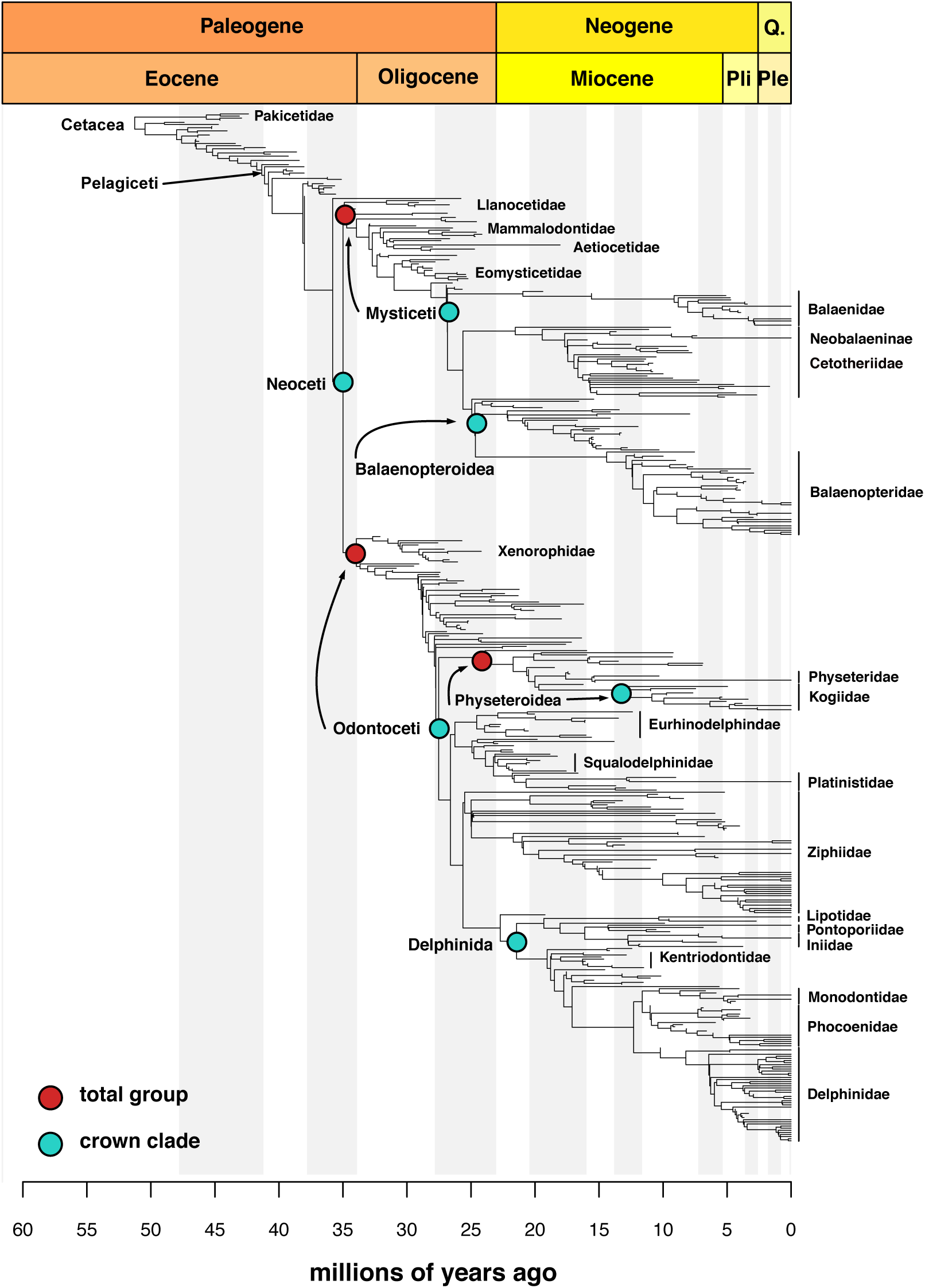
Maximum *a posteriori* chronogram derived from simultaneous Bayesian inference of topology and branch lengths. The strict consensus metatree derived from analysis of species–level OTUs is used as a topology constraint with stratigraphic ages for extinct taxa and *Cytochrome B* sequence data for extant taxa used to help resolve polytomies. Shaded bars correspond to marine stages.

Bayesian estimation of branch lengths on the GENUS and ALL datasets resulted in larger time trees with more fossil taxa but yielded substantially older divergence times for some crown clades than we found for the EXCLUDE dataset. To facilitate comparison of our results to divergence time estimates derived from node–dated trees, we extracted mean ages and their associated 95% HPD intervals for select crown clades and compared them to those inferred by McGowen et al. (2019) using a genomic dataset. These estimates (Table show that node age estimates are relatively consistent between the genomic tree and our EXCLUDE metatree, albeit with the metatree generating slightly younger node ages. Node ages for the GENUS and ALL datasets are, on average, a little older than those in the EXCLUDE tree and more similar to those of the genomic tree. Balaenidae and Delphinidae deviate substantially from this pattern, however, with mean age estimates that are approximately 10 and 7 million years older, respectively, than the EXCLUDE dataset and have 95% HPD intervals that do not overlap.

### Inference of Diversification Dynamics

Similar to analyses based on extant cetaceans alone, we found that net diversification rates are relatively constant through time, but with a rapid increase in mean net diversification rates beginning at approximately 10 Ma for the EXCLUDE MAP chronogram. In contrast with inference from molecular phylogenies, this result arises not only from a moderate increase in speciation rates, but also from a precipitous decline in extinction rates over the same time frame (Figs 6 b,c). These average rates are clearly emergent properties of more complex, clade-specific dynamics. The 95% credible shift set for the EXCLUDE MAP tree contained 483 distinct configurations, with 2 – 6 shifts recovered most often (Table 2). No individual configuration occurred with any meaningful frequency (*f* = 0.055 or less). However, plots of mean per–branch speciation, extinction, and net diversification rates show that elevated net diversification rates in mesoplodont beaked whales, a result not previously identified in molecular phylogenies, result from depressed rates of extinction against a backdrop of already low rates of speciation, while rapid diversification rates in oceanic dolphins result from both elevated speciation and depressed extinction rates (Figs 7a–c).

**Fig. 6.**
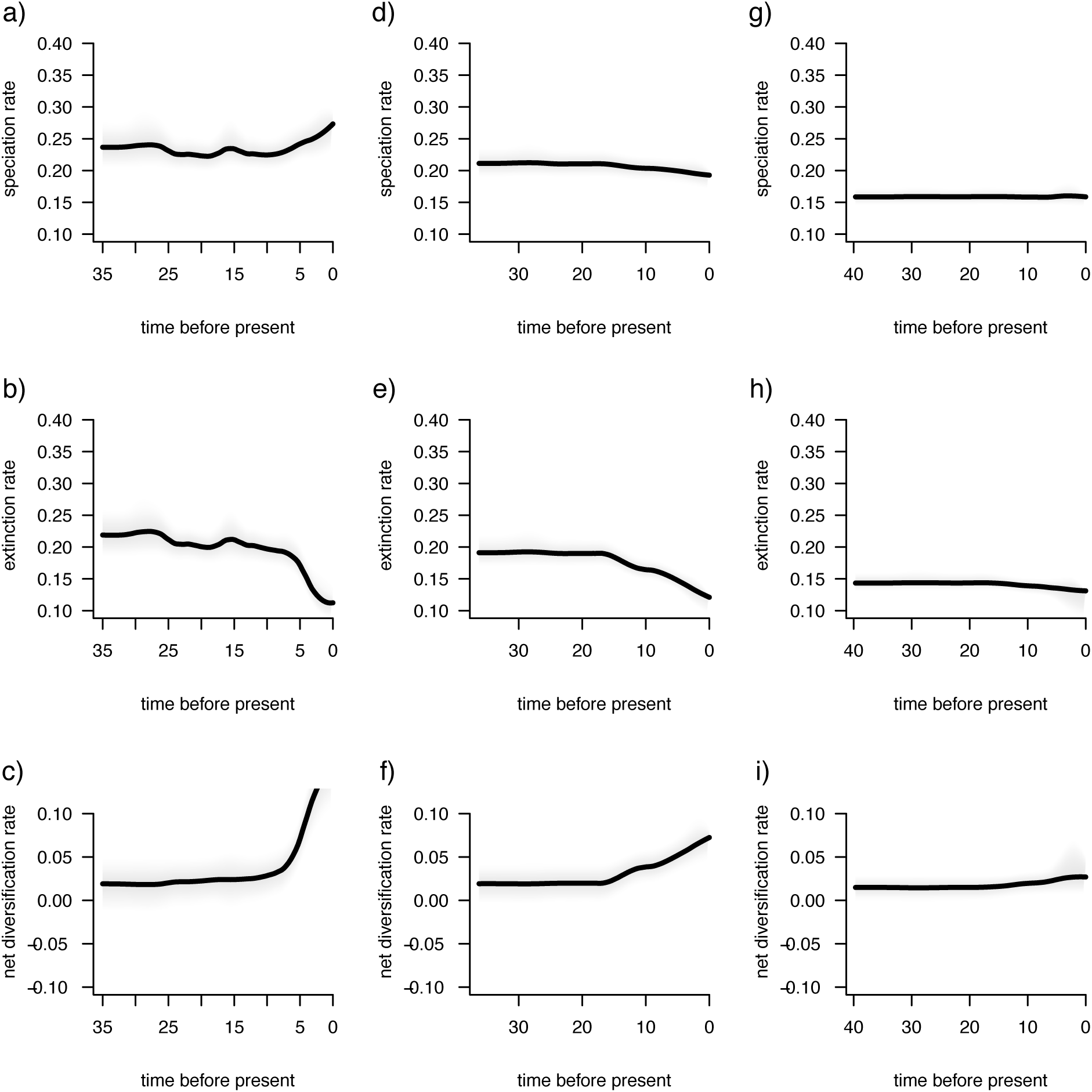
Diversification dynamics through time inferred from the EXCLUDE (a:c), GENUS (d:f), and ALL (g:i) datasets suggest very different dynamics through time. Note that Speciation and Extinction rates are plotted on the same scale as each other, and as in (Rabosky, 2014, figure 9D:E) for extant cetaceans.

**Fig. 7.**
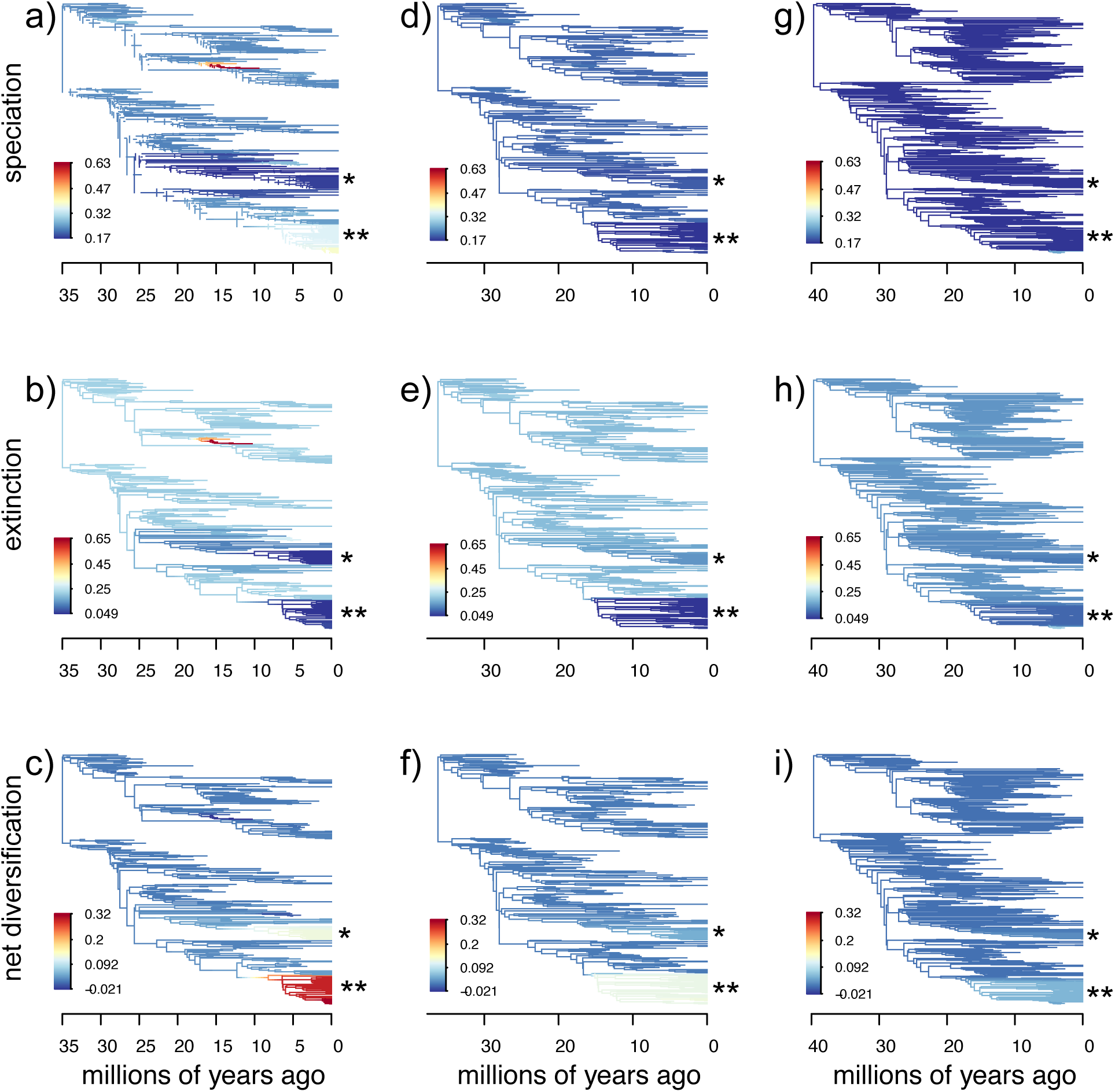
Mean per–branch rates of speciaton, extinction, and net–diversification rates for the EXCLUDE (a:c), GENUS (d:f), and ALL (g:i) datasets. As in Figure 6, the top row is speciation rates, middle row is extinction rates, and bottom row is net diversification rate. The single asterix (*) denotes mesoplodont beaked whales, the double asterices (**) denote oceanic dolphins (Delphinidae). Note that the same scale is used for each plot to enable comparisons of absolute magnitudes of the underlying estimated rates.

Increases in mean net diversification rates towards the present day are more muted in the GENUS chronogram (Figs 6 d–f). Although we recovered similarly declining mean extinction rates as for the EXCLUDE tree, we found no increase in mean speciation rates and, in fact, recovered a slight decline over this time–frame (Figs 6d,e). The number of inferred shifts is much lower for the GENUS dataset (Table 2), and only 83 possible configurations are present in the 95% credible set. Mean per–branch rates (Figs 7d–f) show that the more muted increases in net diversification for mesoplodontid ziphiids and ocean dolphins arise from decreased extinction rates in these clades.

Rate variation is further dampened in the ALL analysis. Here, there is no increase in mean net diversification rate and both mean speciation and extinction rates have remained relatively low and constant, albeit with a very slight increase in speciation and decline in extinction at approximately 10 Ma (Figs. 6g–i). Although a configuration with no shifts was the most frequently sampled (Table 2), 154 alternative configurations are present in the 95% credible set. Mean per–branch rates of speciation and extinction are relatively homogeneous, but with very slight increases in speciation and decreases in extinction rates, leading to slight increases in net diversification in delphinids (Figs. 7g:i).

## Discussion

The ability to infer comprehensively sampled phylogenies of extant higher–level clades has led to novel hypotheses regarding their macroevolutionary dynamics (Smith et al., 2009; Smith and Brown, 2018; Jetz et al., 2012; Zanne et al., 2014; Cooney et al., 2017; Tonini et al., 2016; Jetz and Pyron, 2018; Rabosky et al., 2018; Upham et al., 2019; Varga et al., 2019). However, even a limited amount of data from fossil taxa can overturn well-supported hypotheses derived from analyses of extant taxa only (Finarelli and Flynn, 2006; Albert et al., 2009; Slater et al., 2012; Betancur-R et al., 2015; Meseguer et al., 2014) and it is likely that datasets consisting exclusively or primarily of fossil taxa are needed to test fundamental macroevolutionary hypotheses. The most substantial barrier to implementing such tests has always been the difficulty in assembling robust, time–scaled phylogenies for higher–level clades that contain sufficient numbers of fossil taxa. Our well–resolved cetacean metatrees suggests that meta–analytic phylogenies can provide a useful and compelling way of synthesizing studies of lower–level clades to produce higher–level phylogenetic hypotheses for extinct taxa. Moreover, these trees provide an accessible way of addressing questions of macroevolutionary importance using fossil data and allow for the interrogation of results derived from phylogenies of extant taxa alone.

### Implications for Cetacean Diversification

It has been recognized for some time that estimates of extinction rates derived from molecular phylogenies may be problematic (Rabosky, 2010; Beaulieu and O’Meara, 2015). Empirical studies have found that diversification rates estimated from molecular phylogenies may be congruent with inferences derived from paleontological data but often differ in the underlying estimates of speciation and extinction rates over time (e.g, Simpson et al., 2011; Cantalapiedra et al., 2015; Hagen et al., 2017; Law et al., 2017). The myriad ways in which speciation and extinction rates can vary to produce identical lineage through time plots for phylogenies of extant species was recently emphasized by Louca and Pennell (2020). One of the many implication of their work is that the dimensionality of model space (that is, the number of possible combinations of time–varying speciation and extinction rates) is too large to reliably identify the generating model when only extant species are sampled and that robust inference of speciation and extinction rates through time can only be achieved with densely sampled phylogenies that incorporate extinct and extant lineages.

One of the most striking results to emerge from our diversification analyses is that variation in extinction rates, rather than speciation rates, have played a dominant role in shaping extant cetacean diversity. It is, of course, a mathematical necessity that rates of speciation must increase or rates of extinction decline in order for net diversification rates to increase. But, while many neontologists have (explicitly or implicitly) assumed a dominant role for elevated rates of speciation in driving diversification in exceptionally species–rich clades as a response to increased ecological opportunity (for reviews, see Schluter, 2000; Glor, 2010; Stroud and Losos, 2016; Martin and Richards, 2019), paleontologists have tended to recognize a role for extinction rate variation in facilitating radiations over geologic time–scales (Jablonski et al., 1983; Van Valen, 1985; Labandeira and Sepkoski, 1993; Valentine, 1990). However, the difficulty of inferring extinction from molecular phylogenies means that the effects of extinction rate variation have received little attention in phylogenetic contexts. Here, by incorporating fossil taxa in a phylogenetic framework, we found that mesoplodont beaked whales emerge as a previously unidentified rapid radiation. Despite accounting for 15 of 21 extant species Mead and Brownell Jr (1993), this radiation is not characterized by elevated speciation rates but, rather, by depressed extinction (Fig. 7b). Fossil evidence from diverse taxa has showed that clade–level origination and extinction rates tend to be positively correlated (Stanley, 1979), meaning that clades with a higher instantaneous probability of speciating tend to also have a higher long–term probability of going extinct (higher volatility: Gilinsky, 1994), while clades with low extinction probabilities are more extinction resistant (Valentine, 1990). Recent work has found a strong link between ecological diversity and low volatility across living and extinct clades of marine animals (Knope et al., 2020), suggesting that low extinction at the clade level may arise due to factors such as ecological flexibility. Unfortunately, too little is currently known about mesoplodont ecology to derive reasonable hypotheses to explain their low extinction rates and macroevolutionary success.

Diversification studies based on extant cetacean phylogenies have consistently identified the oceanic dolphins as a rapid radiation due to elevated speciation rates during the past 10 myr (McGowen et al., 2009; Steeman, 2010; Slater et al., 2010; Rabosky et al., 2014). Although we still recover Delphindae as rapid radiation using a phylogeny of extant and extinct cetaceans, we find support for strikingly different underlying dynamics. While there is still evidence for increased speciation rates in Delphinidae (Figs 6a; 7a), their elevated net diversification rates are predominantly driven by dramatically decreased extinction rates relative to other cetaceans (Figs 6b; 7b). One explanation for this finding could be that dolphins are in the early phase of adaptive radiation within an unoccupied adaptive zone, wherein speciation is rapid and extinction 0 due to a lack of competition (Simpson, 1953; Valentine, 1980; Van Valen, 1985). However, Stanley (1990) has argued that, because so few clades break the strong correlation between origination and extinction rates, those that do (“Supertaxa”) likely possess uniquely advantageous combinations of life history traits, such as low dispersal rates combined with large population sizes, compared with related clades. The precise nature of the relationships that arise between traits and speciation / extinction dynamics are complex and mechanism dependent (see, for example, Table 1 in Jablonski, 2008) but it is notable that delphinids are social and ecologically flexible, while their diversification has previously been linked to Plio–Pleistocene changes in ocean currents that resulted in abrupt, localized, soft barriers to gene flow (do Amaral et al., 2018). A greater understanding of the multivariate structure of life history traits with Cetacea (e.g.,. Pianka et al., 2017) may reveal more insights into how Delphinidae has managed to break the speciation – extinction correlation with such dramatic effect.

**Table 1.** Divergence time estimates (mean and 95% HPD intervals) for select crown clades from the genomic study of McGowen et al. (2019) and the three metatree analysis.

| Clade | McGowan et al. (2019) | EXCLUDE | GENUS | ALL |
| --- | --- | --- | --- | --- |
| Neoceti | 36.7 (37.5–36.4) | 36.8 (38.4–35.2) | 37.8 (39.3–35.4) | 40.3 (42.0–38.3) |
| Mysticeti | 25.7 (26.7–25.2) | 23.3 (28.1–24.4) | 26.8 (28.4–24.5) | 28.7 (31.2–26.4) |
| Balaenidae | 10.6 (12.1–9.2) | 6.9 (8.3–5.9) | 17.2 (18.4–15.5) | 17.8 (19.6–16.0) |
| Balaenopteridae | 15.7 (16.9 – 14.7) | 13.0 (16.2–9.7) | 15.9 (18.5–12.3) | 20.5 (22.4–18.4) |
| Odontoceti | 34.1 (34.9–33.7) | 28.0 (26.7–29.5) | 28.6 (29.7–26.8) | 31.9 (33.5–30.2) |
| Physteroidea | 22.4 (24.1–20.6) | 21.0 (23.2–18.4) | 23.1 (24.5–21.2) | 25.0 (27.7–22.4) |
| Delphinida | 25.1 (26.1–24.2) | 21.4 (23.7–19.2) | 22.9 (25.1–20.3) | 23.2 (25.1–21.4) |
| Delphinidae | 12.7 (13.6–11.8) | 8.7 (10.6–6.7) | 15.7 (16.7–13.8) | 17.4 (19.7–15.4) |

**Table 2.**
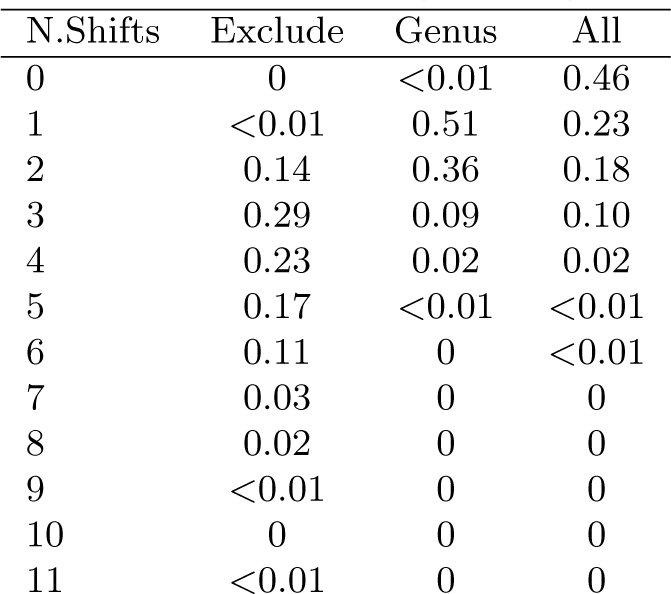
Posterior probabilities for numbers of rate shifts (N. shifts) on the 3 MAP time–scaled metatrees.

### Paleo–Problems and Future Directions

Any phylogenetic hypothesis is only as robust as the data from which it is inferred. Ultimately the onus is on the user to ensure that the data are of sufficient quality and independence that the resulting tree(s) stand up to scrutiny (Bininda-Emonds et al., 2004). By establishing a formalized set of rules for dealing with data re–use, recovery of multiple optimal trees, and the use of OTUs corresponding to different levels of the taxonomic hierarchy, the metatree pipeline (Lloyd et al., 2016) provides an explicit framework for ameliorating some of the criticisms and concerns leveled at earlier MRP supertrees (Springer and de Jong, 2001; Gatesy et al., 2002, 2004; Gatesy and Springer, 2004; Page, 2004). This is not to say that there are no concerns or areas for future improvement with our approach.

MRP supertrees have been criticized on the basis that they can recover unique clades that are not present in the profile of source trees (Wilkinson, 1995; Bininda-Emonds, 2003; Gatesy et al., 2004). Few unusual clades emerge in our strict consensus metatrees, but we do recover a unique Llanocetidae (Fig 5), consisting of *Llanocetus denticrenatus, Mystacodon selenensis, Niparajacetus palmidentis* and ZMT62 an undescribed taxon from New Zealand (Fordyce, 1989). *Mystacodon*’s placement is not a subject of concern; although the taxon was originally described as an earlier diverging mysticete (Lambert et al., 2017; de Muizon et al., 2019) its placement within Llanocetidae is in line with a number of recent studies (e.g., Fordyce and Marx, 2018; Marx et al., 2019; Azucena Solis-Añorve and Gerardo González-Barba and René Hernández-Rivera, 2019). The other two taxa have not been recovered as llanocetids in published sources incorporated here, but their placements can be easily explained. ZMT62 features in a single study, that of Geisler et al. (2017), and is figured (their Fig. 4) as the sister lineage to a clade consisting of Mammalodontidae + Aetiocetidae and ((*Llanocetus*, (Eomysticetidae, crown group mysticetes)). Inspection of the supplementary methods of Geisler et al. (2017) reveals that this topology is derived from an analysis using implied weights (Goloboff, 1993), a procedure that has been shown to increase resolution at the expense of accuracy (Congreve and Lamsdell, 2016), and that the authors’ own analyses using equal weights yield a topology with ZMT62 as the sister taxon to *Llanocetus*, as we also found. The placement of *Niparajacetus* can be equally well explained. The original description of this taxon included a bootstrap consenus tree rooted sequentially by the archaeocete *Zygorhiza* and a selection of odontocetes, in which *Niparajacetus* is recovered in a polytomy with *Coronodon havensteini*, Mammalodontidae, Aetiocetidae, Eomysticetidae and crown group mysticetes, and with Llanocetidae as sister to this clade (Azucena Solis-Añorve and Gerardo González-Barba and René Hernández-Rivera, 2019, their Fig.7). Our reanalysis of the character matrix yields an identical topology with one exception: the odontocete outgroups are nested within mysticetes. Indeed, rerooting the tree on the odontocetes produces a topology in which *Niparajacetus* falls within a monophyletic Llanocetidae, consistent with our metatree results. The recovery of this previously unreported clade in the metatree can therefore be considered to result from careful scrutiny of the input data, rather than a compromise between conflicting relationships in figured topologies that might emerge from a traditional MRP supertree.

It is well understood that failing to account for unsampled taxa can bias inference of diversification dynamics based on molecular phylogenies (Pybus and Harvey, 2000; FitzJohn et al., 2009; Höhna et al., 2011). To overcome this issue, some authors have used random birth–death resolutions, combined with taxonomic constraints, to integrate over all possible placements of unsampled taxa (Kuhn et al., 2011). We used a similar procedure here to include unsampled fossil species by using the Paleobiology Database’s taxonomy as a down–weighted constraint during metatree inference. There are reasons to be concerned that this procedure may introduce a substantial source of error when inferring the placement of unsampled species. Although cetaceans possess one of the most well–curated set of records in the database (e.g., Uhen and Pyenson, 2007), a number of records of uncertain or doubtful status exist that have dramatic impacts on downstream analyses. For example, the Paleobiology Database records the taxon *Balaena dubusi* from the middle Miocene of Belgium (Louwye et al., 2010), in turn implying a minimum age of 15 Ma for the divergence of the sole extant member of the genus *Balaena*, the bowhead *B. mysticetus* from the right whales *Eubalaena*. In our GENUS and ALL analyses, inclusion of this taxon contributes to an increase in the mean age of crown group balaenids from 8.6 Ma in the EXCLUDE analysis to ∼ 18 Ma (Table 1). *B. dubusi* was described by Van Beneden (1872) from a single vertebral column and Steeman (2010) has discussed the many issues surrounding taxa described from the Antwerp faunas, considering many as *nomina dubia*. While the status of *B. dubusi* awaits formal re–assessment, it seems plausible that its assignment to the extant genus *Balaena* is in error. It is likely that similar taxonomic issues influence topology, branch lengths, and subsequent macroevolutionary inference in other parts of the GENUS and ALL trees (Figures 6,7). Notably, the inference of older divergence times for Delphinidae in these “more complete” trees than for the EXCLUDE analysis may provide an explanation for the loss of signal for increased diversification rates during the past 10 Ma. It should be noted that the birth–death polytomy resolution is not without issue in molecular phylogenetics either, resulting in elevated relative extinction rates (*µ* / λ), increased “tippy–ness” and more balanced trees than are found in empirical distributions of trees (Kuhn et al., 2011). The appropriate placement of unsampled extant taxa similarly depends on the accuracy of taxonomic constraints used. However, the fact that fossil taxa are non–contemporaneous means that they potentially exert more influence on divergence time estimates (Soul and Friedman, 2015). The inclusion of unsampled fossil taxa in meta–analytic phylogenies should always be carefully considered and justified and, at least for cetaceans, we recommend that the EXCLUDE trees should be the preferred hypotheses used in downstream analyses. More generally, these results emphasize that uncritical use of paleontological databases in phylogenetic and macroevolutionary research has the potential to produce flawed inferences and every taxon should, ideally, be vetted against the literature to corroborate its status.

A potential criticism of the metatree approach, as applied here, is that the resulting posterior distribution of time–scaled topologies does not explicitly incorporate topological uncertainty derived from the sample of input trees. Previous paleo–supertree studies have attempted to accommodate topological and divergence time uncertainty by first obtaining a subsample of most parsimonius trees and then using paleontological approaches to generate multiple sets of branch lengths per tree (e.g., Clarke et al., 2016; Lloyd et al., 2016). Although some phylogenetic uncertainty is propagated through our BEAST analyses due to the use of the strict consensus metatree as a topological constraint, many nodes were fixed (for the EXCLUDE tree in particular) due to the well–resolved nature of the resulting estimate. Molecular phylogeneticists have employed a divide–and–conquer approach called “backbone–and–patch” (Jetz et al., 2012; Tonini et al., 2016; Jetz and Pyron, 2018; Upham et al., 2019), wherein topologies for densely sampled monophyletic subclades are pasted onto time–scaled higher–level topologies, to obtain a pseudo–posterior distribution of time–scaled topologies that can be used in comparative analyses.

Logistically, such an approach cannot work in paleontological contexts because it would require assumptions of monophyly, which may vary between studies, and appropriate character taxon matrices for both the backbone and patch clades, which are also lacking in most cases. There is some cause to be optimistic that solutions can be found. Akanni et al. (2015) used Markov chain Monte Carlo to sample the posterior distribution of rooted supertree topologies under the exponential error model of Steel and Rodrigo (2008) and found that the approach performed well in terms of topology inference, clade support, and computation time. Efficient approaches for generating time–scaled trees of extinct taxa that also appropriately accommodate topological and branch length uncertainty will require similar Bayesian treatments.

### Conclusions

Metatrees have some key benefits over traditional MRP supertrees that render them ideal for comparative paleobiologists. Complete sampling of taxa from the source data is always achieved, whereas a supertree can suffer when figured source trees collapse non-focal clades (loss of resolution) or key outgroups are excluded (loss of overlap). Additionally, a preferred method of inference (e.g., parsimony, maximum likelihood, or Bayesian inference) can be applied when re-analyzing the source data and a preferred output, such as a complete set of most parsimonious trees, maximum likelihood tree or sample from a Bayesian posterior distribution, can be used for metatree inference. This latter step enables a more realistic inclusion of phylogenetic uncertainty in the resulting composite phylogeny than can be accomplished through the use of published consensus trees (see Bell and Lloyd, 2015, their Figure 4). In other words, metatrees take more information forward from the source data to the synthetic hypothesis than traditional MRP supertrees do, and this tends to lead to better resolved topologies (Lloyd et al., 2016). Most importantly, and as our comparative analyses demonstrate, the ability to generate synthetic phylogenies containing large numbers of extinct taxa allows for the critical assessment of macroevolutionary hypotheses derived from extant taxa alone. Here, we showed that the apparent pulse of increased cetacean diversification during the past 10 myr is driven more by reduced extinction rates than by increased speciation, a pattern long established in the fossil record but almost undetectable using extant species alone. While a supermatrix, with character states coded for every extinct species, remains a compelling standard for morphologists to strive for (Gatesy and Springer, 2004), supertrees realistically provide the most direct and accessible route for generating large phylogenies containing extinct taxa which, as simulations suggest (Slater et al., 2012; Louca and Pennell, 2020) and our results show, are essential for obtaining accurate parameter estimates and model inference from macroevolutionary and macroecological analyses.

## Supporting information

Supplementary Bibliography

## Acknowledgements

We extend our sincere gratitude to Mark D. Uhen for his dedicated entry of cetacean fossil occurrences into the Paleobiology Database. Michael McGowan provided his molecular alignment and discussion of divergence times and the taxonomy of fossil cetaceans. Jon Hill wrote parallelization code for TNT. David Černý, Natalie Cooper, David Jablonski, Felix Marx, Rossy Natale, Mark Uhen, and Anna Wisniewski provided helpful comments and criticisms on early drafts of the manuscript. GTL was supported, in part, by a Researcher Mobility Award from the University of Leeds.

