## Supplementary Bibliography for "A Total-Group Phylogenetic Metatree for Cetacea and the Importance of Fossil Data in Diversification Analyses"

Murakami, M., Shimada, C., Hikia, Y. and Hirano, H., in press. New small phocoenid from the Pliocene of Hokkaido, northern Japan: insight into the growth rate and vertebral evolution of porpoises. *Acta Palaeontologica Polonica*.

Nelson, M. D. and Uhen, M. D., in press. First occurrence of a squalodelphinid (Cetacea, Odontoceti) from the early Miocene of Washington State. *Journal of Vertebrate Paleontology*.

O'Leary, M. A. and Gatesy, J., 2008. Impact of increased character sampling on the phylogeny of Cetartiodactyla (Mammalia): combined analysis including fossils. *Cladistics*, **24**, 397-442.

Paolucci, F., Buono, M. R., Fernandez, M. S., Marx, F. G. and Cuitino, J. I., in press. *Diaphorocetus poucheti* (Cetacea, Odontoceti, Physeteroidea) from Patagonia, Argentina: one of the earliest sperm whales. *Journal of Systematic Palaeontology*.

Peredo, C. M. and Pyenson, N. D., 2018. *Salishicetus meadi*, a new aetiocetid from the late Oligocene of Washington State and implications for feeding transitions in early mysticete evolution. *Royal Society Open Science*, **5**, 172336.

Peredo, C. M. and Uhen, M. D., 2016. A new basal chaeomysticete (Mammalia: Cetacea) from the Late Oligocene Pysht Formation of Washington, U.S.A. *Papers in Palaeontology*, **2**, 533-554.

Peredo, C. M., Pyenson, N. D., Marshall, C. D. and Uhen, M. D., 2018. Tooth loss precedes the origin of baleen in whales. *Current Biology*, **28**, 1-9.

Peredo, C. M., Uhen, M. D. and Nelson, M. D., in press. A new kentriodontid (Cetacea: Odontoceti) from the early Miocene Astoria Formation and a revision of the stem delphinidan family Kentriodontidae. *Journal of Vertebrate Paleontology*.

Post, K., Louwye, S. and Lambert, O., 2017. *Scaldiporia vandokkumi*, a new pontoporiid (Mammalia, Cetacea, Odontoceti) from the Late Miocene to earliest Pliocene of the Westerschelde estuary (The Netherlands). *PeerJ*, **5**, e3991.

Pyenson, N. D., Velez-Juarbe, J., Gutstein, C. S., Little, H., Vigil, D. and O'Dea, A., 2015. *Isthminia panamensis*, a new fossil inioid (Mammalia, Cetacea) from the Chagres Formation of Panama and the evolution of 'river dolphins' in the Americas. *PeerJ*, **3**, 1227.

Racicot, R. A., Boessenecker, R. W., Darroch, S. A. F. and Geisler, J. H., 2019. Evidence for convergent evolution of ultrasonic hearing in toothed whales (Cetacea: Odontoceti). *Biology Letters*, **15**, 20190083.

Ramassamy, B., 2016. Description of a new long-snouted beaked whale from the Late Miocene of Denmark: evolution of suction feeding and sexual dimorphism in the Ziphiidae (Cetacea: Odontoceti). *Zoological Journal of the Linnean Society*, **178**, 381-409.

Sanders, A. E. and Geisler, J. H., in press. A new basal odontocete from the upper Rupelian of South Carolina, U.S.A., with contributions to the systematics of *Xenorophus* and *Mirocetus* (Mammalia, Cetacea). *Journal of Vertebrate Paleontology*.

Shipps, B. K., Peredo, C. M. and Pyenson, N. D., 2019. *Borealodon osedax*, a new stem mysticete (Mammalia, Cetacea) from the Oligocene of Washington State and its implications for fossil whale-fall communities. *Royal Society Open Science*, **6**, 182168.

Solis-Anorve, A., Gonzalez-Barba, G. and Hernandez-Rivera, R., 2019. Description of a new toothed mysticete from the Late Oligocene of San Juan de La Costa, B.C.S., Mexico. *Journal of South American Earth Sciences*, **89**, 337-346.

Spaulding, M., O'Leary, M. A. and Gatesy, J., 2009. Relationships of Cetacea (Artiodactyla) among mammals: increased taxon sampling alters interpretations of key fossils and character evolution. *PLoS One*, **4**, e7062.

Steeman, M. E., 2007. Cladistic analysis and a revised classification of fossil and recent mysticetes. *Zoological Journal of the Linnean Society*, **150**, 875-894.

Tanaka, Y. and Fordyce, R. E., 2014. Fossil dolphin *Otekaikea marplesi* (latest Oligocene, New Zealand) expands the morphological and taxonomic diversity of Oligocene cetaceans. *PLOS One*, **9**, e107972.

Tanaka, Y. and Fordyce, R. E., 2015. A new Oligo-Miocene dolphin from New Zealand: *Otekaikea huata* expands diversity of the early Platanistoidea. *Palaeontologia Electronica*, **18**, 1-71.

Tanaka, Y. and Fordyce, R. E., 2016. Papahu-like fossil dolphin from Kaikoura, New Zealand, helps to fill the Early Miocene gap in the history of Odontoceti. *New Zealand Journal of Geology and Geophysics*, **59**, 551-567.

Tanaka, Y. and Fordyce, R. E., in press. *Awamokoa tokarahi*, a new basal dolphin in the Platanistoidea (late Oligocene, New Zealand). *Journal of Systematic Palaeontology*.

Tanaka, Y. and Watanabe, M., in press. An early and new member of Balaenopteridae from the upper Miocene of Hokkaido, Japan. *Journal of Systematic Palaeontology*.

Tanaka, Y., Ando, T. and Sawamura, H., 2018. A new species of Middle Miocene baleen whale from the Nupinai Group, Hikatagawa Formation of Hokkaido, Japan. *PeerJ*, **6**, e4934.

Theodor, J. M. and Foss, S. E., 2005. Deciduous dentitions of Eocene cebochoerid artiodactyls and cetartiodactyl relationships. *Journal of Mammalian Evolution*, **12**, 161-181.

Thewissen, J. G. M., Williams, E. M., Roe, L. J. and Hussain, S. T., 2001. Skeletons of terrestrial cetaceans and the relationship of whales to artiodactyls. *Nature*, **413**, 277-281.

Thewissen, J. G. M., Cooper, L. N., Clementz, M. T., Bajpai, S. and Tiwari, B. N., 2007. Whales originated from aquatic artiodactyls in the Eocene epoch of India. *Nature*, **450**, 1190-1194.

Tsai, C.-H. and Fordyce, R. E., 2018. A new archaic baleen whale *Toipahautea waitaki* (early Late Oligocene, New Zealand) and the origins of crown Mysticeti. *Royal Society Open Science*, **5**, 172453.

Tsai, C.-H. and Fordyce, R. E., in press. The earliest gulp-feeding mysticete (Cetacea: Mysticeti) from the Oligocene of New Zealand. *Journal of Mammal Evolution*.

Tsai, C.-H. and Fordyce, R. E., in press. Archaic baleen whale from the Kokoamu Greensand: earbones distinguish a new Late Oligocene mysticete (Cetacea: Mysticeti) from New Zealand. *Journal of the Royal Society of New Zealand*.

Uhen, M. D., 1999. New species of protocetid archaeocete whale, *Eocetus wardii* (Mammalia: Cetacea) from the Middle Eocene of North Carolina. *Journal of Paleontology*, **73**, 512-528.

Uhen, M. D., 2004. Form, function, and anatomy of *Dorudon atrox* (Mammalia, Cetacea): an archaeocete from the Middle to Late Eocene of Egypt. *University of Michigan Papers on Paleontology*, **34**, 1-222.

Uhen, M., 2014. New material of *Natchitochia jonesi* and a comparison of the innominata and locomotor capabilities of Protocetidae. *Marine Mammal Science*, **30**, 1029-1066.

Uhen, M. D. and Gingerich, P. D., 2001. New genus of dorudontine archaeocete (Cetacea) from the Middle-to-Late Eocene of South Carolina. *Marine Mammal Science*, **17**, 1-34.

Uhen, M. D., Pyenson, N. D., Devries, T. J., Urbina, M. and Renne, P. R., 2011. New Middle Eocene whales from the Pisco Basin of Peru. *Journal of Paleontology*, **85**, 955-969.

Velez-Juarbe, J., Wood, A. R., De Gracia, C. and Hendy, A. J. W., 2015. Evolutionary patterns among living and fossil kogiid sperm whales: evidence from the Neogene of Central America. *PLOS ONE*, **10**, e0123909.

Velez-Juarbe, J., in press. A new stem odontocete from the late Oligocene Pysht Formation in Washington State, U.S.A. *Journal of Vertebrate Paleontology*.

Viglino, M., Buono, M. R., Fordyce, R. E., Cuitino, J. I. and Fitzgerald, E. M. G., 2019. Anatomy and phylogeny of the large shark-toothed dolphin *Phoberodon arctirostris* Cabrera, 1926 (Cetacea: Odontoceti) from the early Miocene of Patagonia (Argentina). *Zoological Journal of the Linnean Society*, **185**, 511-542.

Viglino, M., Buono, M. R., Gutstein, C. S., Cozzuol, M. A. and Cuitiño, J. I., in press. A new dolphin from the early Miocene of Patagonia, Argentina: insights into the evolution of Platanistoidea in the Southern Hemisphere. *Acta Palaeontologica Polonica*.

Wichura, H., Jacobs, L. L., Lin, A., Polcyn, M. J., Manthi, F. K., Winkler, D. A., Strecker, M. R. and Clemens, M., 2015. A 17-My-old whale constrains onset of uplift and climate change in east Africa. *Proceedings of the National Academy of Sciences, USA*, **112**, 3910-3915.

### References for excluded source data (to recently discovered)

Benites-Palomino, A., Velez-Juarbe, J., Salas-Gismondi, R. and Urbina, M., in press. *Scaphokogia totajpe*, sp. nov., a new bulky-faced pygmy sperm whale (Kogiidae) from the late Miocene of Peru. *Journal of Vertebrate Paleontology*.

Duboys de Lavigerie, G., Bosselaers,, M., Goolaerts, S., Park, T., Lambert, O. and Marx, F. G., in press. New Pliocene right whale from Belgium informs balaenid phylogeny and function. *Journal of Systematic Palaeontology*.

Lambert, O., Auclair, C., Cauxeiro, C., Lopez, M. and Adnet, S., 2018. A close relative of the Amazon river dolphin in marine deposits: a new Iniidae from the late Miocene of Angola. *PeerJ*, **6**, e5556.
