## Supplementary Table for "A Total-Group Phylogenetic Metatree for Cetacea and the Importance of Fossil Data in Diversification Analyses"

**Table S1:** Maximum and minimum dates for first appearances of fossil cetacean taxa used in Bayesian estimation of topology and divergence times. Citations are provided for data that superceded ranges obtained from the Paleobiology Database

| Taxon | Max | Min | Source |
|---|---|---|---|
| *Acrophyseter deinodon* | 7.246 | 5.333 | PBDB |
| *Acrophyseter robustus* | 13.82 | 11.62 | PBDB |
| *Acrophyseter sp.* MUSM 2182 | 6.9 | 6.7 | Lambert et al. (2014) |
| *Aegicetus gehennae* | 38.0 | 33.9 | PBDB |
| *Aegyptocetus tarfa* | 41.3 | 38 | PBDB |
| Aetiocetidae OCPC 1178 | 18.8 | 17.2 | Marx and Fordyce (2015) |
| Aetiocetidae UWBM 82941 | 27.82 | 23.03 | Burke Museum collections record |
| Aetiocetidae UWBM 87135 | 27.82 | 23.03 | Burke Museum collections record |
| *Aetiocetus cotylalveus* | 31 | 28 | Marx and Fordyce (2015) |
| *Aetiocetus polydentatus* | 26.1 | 23.03 | Marx and Fordyce (2015) |
| *Aetiocetus weltoni* | 28.1 | 28 | Marx and Fordyce (2015) |
| aff *Huaridelphis raimondii* MUSM 603 | 20.44 | 15.97 | Lambert et al. (2016) |
| *Africanacetus gracilis* | 5.333 | 3.6 | PBDB |
| *Africanacetus ceratopsis* | 11.60800 | 5.6 | PBDB |
| *Aglaocetus burtinii* | 11.62 | 7.246 | Steeman (2010) |
| *Aglaocetus latifrons* | 15.97 | 11.62 | Steeman (2010) |
| *Aglaocetus longifrons* | 15.97 | 11.62 | Steeman (2010) |
| *Aglaocetus moreni* | 19.8 | 18.2 | Marx and Fordyce (2015) |
| *Aglaocetus patulus* | 14.5 | 13.9 | Marx and Fordyce (2015) |
| *Aglaocetus rotundus* | 11.62 | 7.246 | Steeman (2010) |
| Agorophiidae ChM PV5852 | 27.82 | 23.03 | Boessenecker and Geisler (2018) |
| *Agorophius pygmaeus* | 27.82 | 23.03 | Boessenecker and Geisler (2018) |
| *Albertocetus meffordorum* | 29 | 26.2 | Boessenecker et al. (2017) |
| *Albertocetus sp* ChM PV4834 | 27.82 | 23.03 | Geisler et al. (2011) |
| *Albicetus oxymycterus* | 15.97 | 13.82 | Boersma and Pyenson (2015) |
| *Albireo savagei* | 3.6 | 2.588 | PBDB |
| *Albireo whistleri* | 7.246 | 5.333 | PBDB |
| *Allodelphis pratti* | 22 | 21 | Toshiyuki and Barnes (2016) |
| *Allodelphis woodburnei* | 22 | 21 | Toshiyuki and Barnes (2016) |
| *Ambulocetus natans* | 47.8 | 41.3 | PBDB |
| *Amphicetus later* | 11.62 | 7.246 | PBDB |
| *Anacharsis orbus* | 12.7 | 11.608 | PBDB |
| *Ancalecetus simonsi* | 37.8 | 33.9 | Gingerich and Uhen (1996) |
| *Andrewsiphius sloani* | 47.8 | 44.5 | Bebej (2011) |
| *Anoplonassa forcipata* | 5.3 | 3.5 | PBDB |
| *Aondelphis talen* | 18.3 | 15.97 | Viglino et al. (2018); Bown and Larriestra (1990) |
| *Aporotus recurvirostris* | 15.97 | 11.62 | PBDB |
| *Aporotus dicyrtus* | 11.6 | 2.588 | Bianucci et al. (2016) |
| *Aprixokogia kelloggi* | 5.333 | 3.6 | PBDB |

| | | | |
|---|---|---|---|
| *Araeodelphis natator* | 17 | 16 | Godfrey et al. (2017) |
| *Archaebalaenoptera castriarquati* | 3.6 | 3.1 | Marx and Fordyce (2015) |
| *Archaebalaenoptera liesselensis* | 8.1 | 7.5 | Bisconti et al. (2020) |
| *Archaeodelphis patrius* | 27.2 | 24.5 | Marx and Fordyce (2015) |
| *Archaeophocaena teshioensis* | 7.246 | 5.333 | PBDB |
| *Archaeoziphius microglenoideus* | 15 | 13.2 | PBDB |
| *Archaeschrichtius ruggieroi* | 11.62 | 7.246 | Bisconti and Varola (2006) |
| *Argyrocetus bakersfieldensis* | 28.1 | 23.03 | PBDB |
| *Argyrocetus joaquinensis* | 28.1 | 23.03000 | PBDB |
| *Argyrocetus patagonicus* | 18.3 | 15.97 | PBDB |
| *Arimidelphis sorbinii* | 3.6 | 1.806 | PBDB |
| *Arktocara yakataga* | 28.1 | 23.03 | PBDB |
| *Artiocetus clavis* | 47.8 | 41.3 | PBDB |
| *Ashleycetus planicapitis* | 33.9 | 28.1 | PBDB |
| *Astadelphis gastaldii* | 5.333 | 2.588 | PBDB |
| *Atocetus iquensis* | 13.82 | 11.62 | PBDB |
| *Atocetus nasalis* | 11.62 | 7.246 | PBDB |
| *Atropatenocetus posteocenicus* | 28.1 | 23.03 | PBDB |
| *Attockicetus praecursor* | 47.8 | 41.3 | PBDB |
| *Aulophyseter mediatlanticus* | 11.62 | 7.246 | PBDB |
| *Aulophyseter morricei* | 15.97 | 13.82 | PBDB |
| *Auroracetus bakerae* | 5.333 | 3.6 | PBDB |
| *Australithax intermedia* | 9.1 | 9 | Laime (2013) |
| *Australodelphis mirus* | 5.333 | 3.6 | PBDB |
| *Austrosqualodon trirhizodonta* | 27.3 | 25.2 | PBDB |
| *Awadelphis hirayamai* | 7.246 | 5.333 | PBDB |
| *Awamokoa tokarahi* | 27.3 | 25.2 | PBDB |
| *Babiacetus indicus* | 47.8 | 38 | PBDB |
| *Babiacetus mishrai* | 47.8 | 41.3 | PBDB |
| *Balaena affinis* | 2.588 | 1.806 | PBDB |
| *Balaena montalionis* | 4.5 | 3.9 | Marx and Fordyce (2015) |
| *Balaena pampaea* | 1.806 | 0.781 | PBDB |
| *Balaena ricei* | 4.9 | 4.4 | Marx and Fordyce (2015) |
| *Balaena arcuata* | 16.5 | 15.1 | Louwye et al. (2010) |
| *Balaena dubusi* | 16.5 | 15.1 | Louwye et al. (2010) |
| *Balaena macrocephalus* | 15.97 | 13.82 | PBDB |
| *Balaena sp* SDSNH 43880 | 5.3 | 2.58 | PBDB |
| *Balaenella brachyrhynus* | 5.0 | 4.4 | Marx and Fordyce (2015) |
| Balaenidae OU 22224 | 27.3 | 26 | Marx and Fordyce (2015) |
| *Balaenoptera bertae* | 3.4 | 2.5 | Marx and Fordyce (2015) |
| *Balaenoptera davidsonii* | 3.6 | 2.588 | PBDB |
| *Balaenoptera siberi* | 8 | 7 | Marx and Fordyce (2015) |
| *Balaenoptera taiwanica* | 5.333 | 3.6 | PBDB |
| *Balaenoptera cephalus* | 15.97 | 13.82 | PBDB |
| *Balaenoptera cuvieri* | 5.3 | 3.0 | Marx and Fordyce (2015) |
| *Balaenoptera sursiplana* | 15.97 | 13.82 | PBDB |
| *Balaenopteridae aka portisi* | 3.6 | 2.588 | Bisconti et al. (2020) |
| Balaenopteridae EGAPA MPTAM 207.13307 | 3.6 | 2.588 | Bisconti and Varola (2006) |
| Balaenopteridae IPMM 40063 | 5.333 | 3.6 | Bisconti et al. (2020) |
| Balaenopteridae MHNL 1610 | 11.608 | 5.332 | Bisconti and Varola (2006) |
| Balaenopteridae MHNL 1613 | 3.6 | 1.806 | Bisconti and Varola (2006) |
| Balaenopteridae NMNZ MM001630 | 5.3 | 3.6 | Bisconti and Varola (2006) |
| Balaenopteridae NMR 999100007096 | 3.6 | 2.588 | Bisconti and Varola (2006) |
| Balaenopteridae RBINS M.2231 | 5.333 | 3.6 | Bisconti et al. (2020) |
| Balaenopteridae RBINS M 2315 | 5.333 | 3.6 | Bisconti et al. (2020) |
| Balaenopteridae RBINS Unreferred | 5.332 | 3.6 | Bisconti and Varola (2006) |
| Balaenopteridae SAMPQL 55001 | 11.63 | 5.33 | Bisconti et al. (2020) |
| Balaenopteridae UT PU13842 /5 | 5.333 | 3.6 | Bisconti et al. (2020) |
| *Balaenotus insignis* | 5.333 | 1.806 | PBDB |

| | | | |
|---|---|---|---|
| *Balaenotus orcianensis* | 3.6 | 2.588 | PBDB |
| *Balaenula astensis* | 4 | 3 | Marx and Fordyce (2015) |
| *Balaenula balaenopsis* | 4 | 2.5 | Bisconti (2003) |
| *Balaenula forsythmajori* | 3.6 | 2.588 | PBDB |
| *Balaenula sp* HUES 10003 | 5.332 | 3.6 | PBDB |
| *Balaenula sp* SMAC 1309 | 6.4 | 4.1 | PBDB |
| Basilosauridae MUSM 1443 | 40.4 | 37.2 | PBDB |
| *Basilosaurus cetoides* | 38 | 33.9 | PBDB |
| *Basilosaurus drazindai* | 41.3 | 38 | PBDB |
| *Basilosaurus isis* | 38 | 33.9 | PBDB |
| *Basiloterus hussaini* | 41.3 | 38 | PBDB |
| *Basilotritus uheni* | 47.8 | 38 | PBDB |
| *Basilotritus wardii* | 41.3 | 38 | PBDB |
| *Basilotritus sp* GMTSNUK 15 | 41.2 | 37.8 | PBDB |
| *Belemnoziphius compressus* | 3.6 | 1.806 | PBDB |
| *Belemnoziphius prorops* | 5.333 | 3.6 | PBDB |
| *Belonodelphis peruanus* | 13.82 | 11.62 | PBDB |
| *Beneziphius cetariensis* | 7.246 | 3.6 | PBDB |
| *Beneziphius brevirostris* | 13.2 | 12.8 | PBDB |
| *Bohaskaia monodontoides* | 5.333 | 3.6 | PBDB |
| *Borealodon osedax* | 30.6 | 28.3 | Shipps et al. (2019) |
| *Brabocetus gigaseorum* | 5.333 | 3.6 | PBDB |
| *Brachydelphis jahuayensis* | 9.63 | 9.38 | Lambert and De Muizon (2013) |
| *Brachydelphis mazeasi* | 13.82 | 9.63 | Lambert and De Muizon (2013) |
| *Brandtocetus chongulek* | 11.2 | 9.6 | Marx and Fordyce (2015) |
| *Brujadelphis ankylorostris* | 13.82 | 11.62 | PBDB |
| *Brygmophyseter shigensis* | 15.97 | 13.82 | PBDB |
| *Carolinacetus gingerichi* | 41.3 | 38 | PBDB |
| *Casatia thermophila* | 5.08 | 4.52 | Bianucci et al. (2019) |
| *Caviziphius altirostris* | 15.97 | 5.33 | PBDB |
| *Cephalotropis coronatus* | 11.7 | 8.5 | Marx and Fordyce (2015) |
| Cetacea ZMT 62 | 33.8 | 28 | Fordyce (1989) |
| *Ceterhinops longifrons* | 29 | 24.7 | PBDB |
| *Cetorhynchus atavus* | 11.62 | 7.246 | PBDB |
| *Cetorhynchus christoli* | 15.97 | 11.63 | PBDB |
| Cetotheriidae ZIRM V28 1 | 15.97 | 11.63 | PBDB |
| *Cetotheriophanes capellinii* | 3.6 | 2.58 | PBDB |
| *Cetotheriopsis lintianus* | 26.84 | 23.19 | PBDB |
| *Cetotherium capellinii* | 5.333 | 2.588 | PBDB |
| *Cetotherium furlongi* | 20.44 | 15.97 | PBDB |
| *Cetotherium parvum* | 15.97 | 13.82 | PBDB |
| *Cetotherium rathkii* | 11.2 | 9.6 | Marx and Fordyce (2015) |
| *Cetotherium riabinini* | 11.2 | 9.6 | Marx and Fordyce (2015) |
| *Cetotherium crassangulum* | 5.33 | 2.58 | PBDB |
| *Cetotherium megalophysum* | 11.7 | 10 | Marx and Fordyce (2015) |
| *Cetotherium polyporum* | 4.9 | 2.8 | PBDB |
| Chaeomysticeti OU 22705 | 21.7 | 20.5 | Marx and Fordyce (2015) |
| Chaeomysticeti ZMT 67 | 21.7 | 20.5 | Marx and Fordyce (2015) |
| *Chavinziphius maxillocristatus* | 6.93 | 6.71 | PBDB |
| *Chilcacetus cavirhinus* | 20.44 | 15.97 | PBDB |
| *Chimuziphius coloradensis* | 8.9 | 7.249 | PBDB |
| *Chonecetus sookensis* | 24.8 | 24.1 | Marx and Fordyce (2015) |
| *Chonecetus tomitai* | 28.1 | 23.03 | PBDB |
| *Choneziphius chonops* | 5.333 | 3.6 | PBDB |
| *Choneziphius leidyi* | 6.1 | 4.4 | PBDB |
| *Choneziphius liops* | 5.333 | 3.6 | PBDB |
| *Choneziphius planirostris* | 9.5 | 7.5 | PBDB |
| *Choneziphius trachops* | 5.333 | 3.6 | PBDB |
| *Choneziphius macrops* | 29 | 24.7 | PBDB |

| | | | |
|---|---|---|---|
| *Chrysocetus fouadassii* | 41.3 | 38 | PBDB |
| *Chrysocetus healyorum* | 38 | 33.9 | PBDB |
| *Ciuciulea davidi* | 13.82 | 12.65 | PBDB |
| *Cophocetus oregonensis* | 20.44 | 15.97 | PBDB |
| *Coronodon havensteini* | 33.9 | 28.1 | PBDB |
| *Cotylocara macei* | 28.1 | 23.03 | PBDB |
| *Crenatocetus rayi* | 47.8 | 41.2 | PBDB |
| *Cynthiacetus maxwelli* | 41.3 | 33.9 | PBDB |
| *Cynthiacetus peruvianus* | 38 | 33.9 | PBDB |
| *Dagonodum mojnum* | 11.62 | 7.246 | PBDB |
| *Dalanistes ahmedi* | 47.8 | 41.3 | PBDB |
| *Dalpiazina ombonii* | 23.03 | 15.97 | PBDB |
| *Delphinapterus brocchii* | 3.6 | 2.588 | PBDB |
| *Delphinavus newhalli* | 13.6 | 10.3 | PBDB |
| *Delphinodon carniolicus* | 15.97 | 13.82 | PBDB |
| *Delphinodon dividum* | 20.44 | 15.97 | PBDB |
| Delphinoidea NMV 72 | 23.03 | 15.97 | PBDB |
| *Delphinus delannoy* | 13.82 | 11.62 | PBDB |
| *Delphinus domeykoi* | 5.333 | 3.6 | PBDB |
| *Delphinus rikuzenensis* | 5.3 | 3.6 | PBDB |
| *Denebola brachycephala* | 7.246 | 5.333 | PBDB |
| *Dhedacetus hyaeni* | 47.8 | 41.3 | PBDB |
| *Diaphorocetus poucheti* | 23.03 | 20.44 | PBDB |
| *Dilophodelphis fordycei* | 20.44 | 15.97 | PBDB |
| *Diorocetus chichibuensis* | 16.4 | 15.1 | Marx and Fordyce (2015) |
| *Diorocetus hiatus* | 14.5 | 13.9 | Marx and Fordyce (2015) |
| *Diorocetus shobarensis* | 17 | 14.9 | Marx and Fordyce (2015) |
| *Diunatans luctoretemergo* | 5 | 4.4 | Marx and Fordyce (2015) |
| *Dorudon atrox* | 38 | 33.9 | PBDB |
| *Dorudon serratus* | 38 | 33.9 | PBDB |
| *Echovenator sandersi* | 27 | 24 | Churchill et al. (2016) |
| *Ediscetus osbornei* | 28.75 | 28.43 | Albright III et al. (2018) |
| *Eobalaenoptera harrisoni* | 13.82 | 11.63 | PBDB |
| *Eocetus schweinfurthi* | 41.3 | 38 | PBDB |
| *Eodelphinus kabatensis* | 11.62 | 7.246 | PBDB |
| *Eomysticetus carolinensis* | 24.7 | 23.5 | PBDB |
| *Eomysticetus whitmorei* | 28.1 | 26.8 | Marx and Fordyce (2015) |
| *Eoplatanista gresalensis* | 23.03 | 20.44 | PBDB |
| *Eoplatanista italica* | 23.03 | 20.44 | PBDB |
| *Eoplatanista miocaenus* | 23.03 | 20.44 | PBDB |
| *Eoplatanista taurinensis* | 20.44 | 15.97 | PBDB |
| *Eosqualodon langewieschei* | 28.1 | 23.03 | PBDB |
| *Eosqualodon latirostris* | 23.03 | 20.44 | PBDB |
| Eschrichtiidae SDSNH 90517 | 5.3 | 2.58 | PBDB |
| *Eschrichtioides gastaldii* | 4 | 3 | Marx and Fordyce (2015) |
| *Eschrichtius akishimaensis* | 1.95 | 1.77 | PBDB |
| *Etruridelphis giulii* | 5.333 | 1.806 | PBDB |
| *Eubalaena ianitrix* | 3.2 | 2.8 | PBDB |
| *Eubalaena shinshuensis* | 6.1 | 5.9 | Marx and Fordyce (2015) |
| *Eubalaena* sp MASCC Unreferred | 3.5 | 3.3 | Bisconti (2002) |
| *Eucetotherium helmersenii* | 12.7 | 11.608 | PBDB |
| *Eudelphis mortezelensis* | 16.5 | 15.1 | PBDB |
| *Eurhinodelphis cocheteuxi* | 16.5 | 15.1 | PBDB |
| *Eurhinodelphis longirostris* | 16.5 | 15.1 | PBDB |
| *Ferecetotherium kelloggi* | 28.1 | 23.03 | PBDB |
| *Fragilicetus velponi* | 5.333 | 3.6 | PBDB |
| *Fucaia goedertorum* | 28.1 | 26.5 | Marx and Fordyce (2015) |
| *Fucaia buelli* | 33.2 | 31 | Marx and Fordyce (2015) |
| *Gandakasia potens* | 47.8 | 41.3 | PBDB |

| | | | |
|---|---|---|---|
| *Gaviacetus razai* | 47.8 | 41.3 | PBDB |
| *Georgiacetus vogtlensis* | 47.8 | 38 | PBDB |
| *Globicephala etruriae* | 3.6 | 2.588 | PBDB |
| *Globicephala uncidens* | 2.588 | 1.806 | PBDB |
| *Globicetus hiberus* | 6.1 | 4.4 | PBDB |
| *Goedertius oregonensis* | 23.03 | 20.44 | PBDB |
| *Goniodelphis hudsoni* | 13.82 | 11.62 | PBDB |
| *Graamocetus longicollis* | 11.62 | 7.246 | PBDB |
| *Gricetoides aurorae* | 4.9 | 3.9 | Marx and Fordyce (2015) |
| *Haborodelphis japonicus* | 5.33 | 3.6 | Bianucci et al. (2019) |
| *Haborophocoena minutus* | 5.333 | 3.6 | PBDB |
| *Haborophocoena toyoshimai* | 5.333 | 3.6 | PBDB |
| *Hadrodelphis calvertense* | 15.97 | 13.82 | PBDB |
| *Hadrodelphis poseidon* | 15.97 | 13.82 | PBDB |
| *Halicetus ignotus* | 13.82 | 11.62 | PBDB |
| *Hemisyntrachelus cortesii* | 5.333 | 2.588 | PBDB |
| *Hemisyntrachelus oligodon* | 7.246 | 5.333 | PBDB |
| *Herentalia nigra* | 11.62 | 7.246 | PBDB |
| *Herpetocetus bramblei* | 6.4 | 4.9 | Marx and Fordyce (2015) |
| *Herpetocetus morrowi* | 3.5 | 2.5 | Marx and Fordyce (2015) |
| *Herpetocetus scaldiensis* | 16.5 | 15.1 | Louwye et al. (2010) |
| *Herpetocetus sendaicus* | 5.333 | 3.6 | PBDB |
| *Herpetocetus transatlanticus* | 4.9 | 4.4 | Marx and Fordyce (2015) |
| *Herpetocetus* sp NMNS PV19540 | 5.332 | 3.6 | PBDB |
| *Herpetocetus* sp (NMNS PV19540, USNM 362901) | 5.333 | 3.6 | El Adli et al. (2014) |
| *Herpetocetus* sp VMW 65 | 2.1 | 0.7 | PBDB |
| *Heterocetus brevifrons* | 16.5 | 15.1 | Louwye et al. (2010) |
| *Heterocetus guiscardii* | 5.333 | 3.6 | PBDB |
| *Heterocetus scriptus* | 11.62 | 7.246 | PBDB |
| *Heterocetus major* | 5.33 | 2.588 | PBDB |
| *Heterodelphis croatica* | 12.7 | 11.608 | PBDB |
| *Heterodelphis klinderi* | 11.62 | 7.246 | PBDB |
| *Heterodelphis leiodontus* | 13.82 | 11.62 | PBDB |
| *Hibacetus hirosei* | 20.44 | 15.97 | PBDB |
| *Himalayacetus subathuensis* | 54 | 53 | Bajpai and Gingerich (1998) |
| *Hoplocetus borgerhoutensis* | 11.62 | 1.806 | PBDB |
| *Hoplocetus crassidens* | 20.44 | 15.97 | PBDB |
| *Hoplocetus ritzi* | 13.82 | 11.62 | PBDB |
| *Hoplocetus curvidens* | 5.333 | 3.6 | PBDB |
| *Horopeta umarere* | 27.3 | 25.2 | PBDB |
| *Huaridelphis raimondii* | 20.44 | 15.97 | PBDB |
| *Ichthyolestes pinfoldi* | 47.8 | 41.3 | PBDB |
| *Idiocetus guicciardinii* | 3.2 | 2.588 | PBDB |
| *Idiophyseter merriami* | 15.97 | 13.82 | PBDB |
| *Idiorophus bolzanensis* | 21.73 | 20.43 | Hampe (2006) |
| *Idiorophus patagonicus* | 20.44 | 15.97 | PBDB |
| *Ihlengesi saldanhae* | 16 | 5.3 | Bianucci et al. (2007) |
| *Imerocetus karaganicus* | 11.62 | 7.246 | PBDB |
| *Imerodelphis thabagarii* | 13.82 | 11.62 | PBDB |
| *Imocetus piscatus* | 7.246 | 3.6 | PBDB |
| *Incacetus broggii* | 23.03 | 20.44 | PBDB |
| *Incakujira anillodefuego* | 11.62 | 7.246 | PBDB |
| *Indocetus ramani* | 47.8 | 41.3 | PBDB |
| *Inermorostrum xenops* | 33.9 | 28.1 | PBDB |
| *Iniopsis caucasica* | 28.1 | 23.03 | PBDB |
| *Inticetus vertizi* | 20.44 | 15.97 | PBDB |
| *Isanacetus laticephalus* | 17.5 | 16 | Marx and Fordyce (2015) |
| *Ischyrorhynchus vanbenedeni* | 11.62 | 7.246 | PBDB |
| *Isocetus depauwi* | 15.97 | 11.62 | PBDB |

| | | | |
|---|---|---|---|
| *Isthminia panamensis* | 7.246 | 5.333 | PBDB |
| *Izikoziphius angustus* | 16 | 5.3 | Bianucci et al. (2007) |
| *Izikoziphius rossi* | 16 | 5.3 | Bianucci et al. (2007) |
| *Janjucetus hunderi* | 28.1 | 25.6 | Marx and Fordyce (2015) |
| *Joumocetus shimizui* | 11.3 | 11.0 | Marx and Fordyce (2015) |
| *Kampholophos serrulus* | 13.82 | 11.62 | PBDB |
| *Kekenodon onamata* | 27.3 | 25.2 | PBDB |
| *Kentriodon diusinus* | 15.97 | 13.82 | PBDB |
| *Kentriodon fuchsii* | 15.97 | 13.82 | PBDB |
| *Kentriodon hoepfneri* | 13.82 | 11.62 | PBDB |
| *Kentriodon obscurus* | 15.97 | 13.82 | PBDB |
| *Kentriodon pernix* | 20.44 | 13.82 | PBDB |
| *Kentriodon schneideri* | 20.43 | 13.82 | Kazár and Hampe (2014) |
| *Kentriodon hobetsu* | 15.97 | 11.63 | PBDB |
| *Kentriodon nakajimai* | 11.87 | 11.25 | Kimura and Hasegawa (2019) |
| *Kharodacetus sahnii* | 47.8 | 41.3 | PBDB |
| *Khoikhoicetus agulhasis* | 16 | 5.3 | Bianucci et al. (2007) |
| *Khoikhoicetus kergueleni* | 16 | 5.3 | Bianucci et al. (2007) |
| *Kogia pusilla* | 3.6 | 2.588 | PBDB |
| *Kogia* sp MSNTUP I13798 | 5.333 | 2.58 | PBDB |
| Kogiinae ChM VP4994 | 5.333 | 2.58 | PBDB |
| *Kogiopsis floridana* | 13.82 | 11.62 | PBDB |
| *Koristocetus pescei* | 11.62 | 7.246 | PBDB |
| *Kurdalagonus maicopicum* | 13.82 | 11.608 | PBDB |
| *Kurdalagonus mchedlidzei* | 12.1 | 11.2 | Marx and Fordyce (2015) |
| *Kutchicetus minimus* | 47.8 | 41.3 | PBDB |
| *Kwanzacetus khoisani* | 11.62 | 5.33 | Lambert et al. (2018a) |
| *Lagenorhynchus harmatuki* | 5.333 | 3.6 | PBDB |
| *Lamprolithax annectens* | 15.97 | 13.82 | PBDB |
| *Lamprolithax simulans* | 15.97 | 13.82 | PBDB |
| *Leptodelphis stavropolitanus* | 12.7 | 11.608 | PBDB |
| *Liolithax kernensis* | 15.97 | 13.82 | PBDB |
| *Liolithax pappus* | 15.97 | 13.82 | PBDB |
| *Lissodelphis fockii* | 13.82 | 11.608 | PBDB |
| *Lissodelphis nordmanni* | 12.7 | 11.608 | PBDB |
| *Livyatan melvillei* | 9.9 | 8.9 | PBDB |
| *Llanocetus denticrenatus* | 34.2 | 34 | Marx and Fordyce (2015) |
| *Lomacetus ginsburgi* | 11.62 | 7.246 | PBDB |
| *Lonchodelphis occiduus* | 11.6 | 5.3 | PBDB |
| *Lophocetus calvertensis* | 11.62 | 7.246 | PBDB |
| *Lophocetus repenningi* | 11.62 | 7.246 | PBDB |
| *Loxolithax sinuosa* | 15.97 | 13.82 | PBDB |
| *Macrodelphinus kelloggi* | 28.1 | 23.03 | PBDB |
| *Macrokentriodon morani* | 13.82 | 11.62 | PBDB |
| *Macrosqualodelphis ukupachai* | 18.86 | 18.7 | PBDB |
| *Maiabalaena nesbittae* | 33.7 | 30.6 | Prothero et al. (2001); Peredo et al. (2018) |
| *Maiacetus inuus* | 47.8 | 41.3 | PBDB |
| *Makaracetus bidens* | 47.8 | 41.3 | PBDB |
| *Mammalodon colliveri* | 25.7 | 23.9 | Marx and Fordyce (2015) |
| *Mammalodon hakataramea* | 27.3 | 26 | Marx and Fordyce (2015) |
| *Masracetus markgrafi* | 38 | 33.9 | PBDB |
| *Matapanui waihao* | 28.1 | 27.3 | PBDB |
| *Mauicetus parki* | 25.2 | 23 | Marx and Fordyce (2015) |
| *Medocinia tetragorhina* | 20.44 | 15.97 | PBDB |
| *Megaptera hubachi* | 5.33 | 3.6 | PBDB |
| *Meherrinia isoni* | 7.246 | 5.333 | PBDB |
| *Mesocetus agrami* | 12.7 | 11.608 | PBDB |
| *Mesocetus aquitanicus* | 20.44 | 15.97 | PBDB |
| *Mesocetus brachyspondylus* | 15.97 | 13.82 | PBDB |

| | | | |
|---|---|---|---|
| *Mesocetus hungaricus* | 13.82 | 11.1 | PBDB |
| *Mesocetus longirostris* | 16.5 | 15.1 | Louwye et al. (2010) |
| *Mesocetus siphunculus* | 20.44 | 15.97 | PBDB |
| *Mesoplodon longirostris* | 5.333 | 1.806 | PBDB |
| *Mesoplodon posti* | 4.86 | 3.9 | PBDB |
| *Mesoplodon tumidirostris* | 5.333 | 3.6 | PBDB |
| *Mesoplodon slangkopi* | 16 | 5.3 | Bianucci et al. (2007) |
| *Messapicetus gregarius* | 9.1 | 8.1 | PBDB |
| *Messapicetus longirostris* | 10.5 | 8.14 | PBDB |
| *Metasqualodon symmetricus* | 28.1 | 23.03 | PBDB |
| *Metopocetus durinasus* | 11.7 | 10.00 | Marx and Fordyce (2015) |
| *Metopocetus hunteri* | 11.62 | 7.246 | PBDB |
| *Microberardius africanus* | 16 | 5.3 | Bianucci et al. (2007) |
| *Microcetus ambiguus* | 28.1 | 23.03 | PBDB |
| *Microcetus sharkovi* | 28.1 | 23.03 | PBDB |
| *Micromysticetus rothauseni* | 29.6 | 28.1 | Marx and Fordyce (2015) |
| *Micromysticetus tobieni* | 28.73 | 26.57 | PBDB |
| *Microphocaena podolica* | 12.7 | 11.608 | PBDB |
| *Microzeuglodon caucasicum* | 41.2 | 33.9 | PBDB |
| *Miobalaenoptera numataensis* | 6.8 | 6.5 | Tanaka and Watanabe (2019) |
| *Miocaperea pulchra* | 7.5 | 7.3 | Marx and Fordyce (2015) |
| *Miodelphis californicus* | 28.1 | 23.03 | PBDB |
| *Miokogia elongatus* | 20.44 | 15.97 | PBDB |
| *Miophocaena nishinoi* | 7.246 | 5.333 | PBDB |
| *Mirocetus riabinini* | 33.9 | 28.1 | PBDB |
| *Mithridatocetus adygeicus* | 11.62 | 7.246 | PBDB |
| *Mithridatocetus eichwaldi* | 11.62 | 7.246 | PBDB |
| *Mixocetus elysius* | 11.62 | 7.246 | PBDB |
| Monodontidae IRSNB M 1922 | 5.333 | 3.6 | Bianucci et al. (2019) |
| *Morawanocetus yabukii* | 26.1 | 23.3 | Marx and Fordyce (2015) |
| *Morenocetus parvus* | 19.8 | 18.2 | Marx and Fordyce (2015) |
| *Mycteriacetus bellunensis* | 23.03 | 20.44 | PBDB |
| *Mystacodon selenensis* | 38 | 33.9 | PBDB |
| Mysticeti ChM PV4745 | 29.6 | 28.1 | Marx and Fordyce (2015) |
| Mysticeti ChM PV5720 | 28.4 | 23.03 | PBDB |
| Mysticeti OU GS10897 | 33 | 32 | Marx and Fordyce (2015) |
| Mysticeti USNM 314627 | 33.7 | 30.6 | PBDB |
| *Nalacetus ratimitus* | 47.8 | 41.3 | PBDB |
| *Nannocetus eremus* | 11.62 | 7.246 | PBDB |
| *Nannolithax gracilis* | 15.97 | 13.82 | PBDB |
| *Nanokogia isthmia* | 11.62 | 7.246 | PBDB |
| *Natchitochia jonesi* | 41.3 | 38 | PBDB |
| *Nazcacetus urbinai* | 7.55 | 7.3 | PBDB |
| *Nehalaennia devossi* | 8.7 | 8.1 | Bisconti et al. (2020) |
| *Nenga meganasalis* | 16 | 5.3 | Bianucci et al. (2007) |
| *Neosqualodon assenzae* | 20.44 | 15.97 | PBDB |
| *Neosqualodon gastaldii* | 20.44 | 15.97 | PBDB |
| *Ninjadelphis ujiharai* | 20.44 | 15.97 | PBDB |
| *Ninoziphius platyrostris* | 5.9 | 3.9 | PBDB |
| *Niparajacetus palmadentis* | 28.1 | 23.03 | Solis-Añorve et al. (2019) |
| *Norrisanima miocaena* | 7.6 | 7.3 | Marx and Fordyce (2015); Leslie et al. (2019) |
| *Notiocetus romerianus* | 1.806 | 0.781 | PBDB |
| *Notiocetus platensis* | 2.588 | 0.0117 | PBDB |
| *Notocetus vanbenedeni* | 23.03 | 20.44 | PBDB |
| *Notoziphius bruneti* | 11.62 | 7.246 | PBDB |
| *Numataphocoena yamashitai* | 5.333 | 3.6 | PBDB |
| *Ocucajea picklingi* | 41.3 | 38 | PBDB |
| *Odobenocetops leptodon* | 7.246 | 5.333 | PBDB |
| *Odobenocetops peruvianus* | 7.246 | 5.333 | PBDB |

| | | | |
|---|---|---|---|
| Odontoceti CCNHM 1000 | 30.5 | 26.5 | Racicot et al. (2019) |
| Odontoceti ChM PV2764 | 28.4 | 23.03 | PBDB |
| Odontoceti ChM PV4178 | 33.9 | 28.1 | PBDB |
| Odontoceti ChM PV4802 | 28.4 | 23.03 | PBDB |
| *Oedolithax mira* | 15.97 | 13.82 | PBDB |
| *Oligodelphis azerbajdzanicus* | 28.1 | 23.03 | PBDB |
| *Olympicetus avitus* | 30.5 | 26.5 | PBDB |
| *Orcinus citoniensis* | 3.6 | 1.806 | PBDB |
| *Orcinus paleorca* | 1.806 | 0.781 | PBDB |
| *Orycterocetus cornutidens* | 5.333 | 3.6 | PBDB |
| *Orycterocetus crocodilinus* | 20.44 | 7.246 | PBDB |
| *Orycterocetus quadratidens* | 5.333 | 3.6 | PBDB |
| *Orycterocetus* sp MAUL 29/1 | 18 | 6 | Bianucci et al. (2004) |
| *Otekaikea huata* | 27.3 | 25.2 | PBDB |
| *Otekaikea marplesi* | 25.2 | 21.7 | PBDB |
| *Otradnocetus* sp VSEGEI 2401 | 11.608 | 7.246 | PBDB |
| *Otradnocetus* virodovi | 15.97 | 11.608 | PBDB |
| *Pachyacanthus suessii* | 15.97 | 11.608 | PBDB |
| *Pakicetus attocki* | 47.8 | 41.3 | PBDB |
| *Pakicetus calcis* | 47.8 | 41.3 | PBDB |
| *Pakicetus chittas* | 47.8 | 41.3 | PBDB |
| *Pakicetus inachus* | 47.8 | 41.3 | PBDB |
| *Palaeobalaena bergi* | 15.97 | 13.82 | PBDB |
| *Palaeophocaena andrussowi* | 13.82 | 11.62 | PBDB |
| *Papahu taitapu* | 21.7 | 19 | PBDB |
| *Papahu* sp ZMT73 | 23 | 18.7 | PBDB |
| *Pappocetus lugardi* | 41.3 | 38 | PBDB |
| *Parabalaenoptera baulinensis* | 7.6 | 6.7 | Marx and Fordyce (2015) |
| *Parapontoporia pacifica* | 7.246 | 5.333 | PBDB |
| *Parapontoporia sternbergi* | 3.6 | 2.588 | PBDB |
| *Parapontoporia wilsoni* | 11.62 | 7.246 | PBDB |
| *Parietobalaena affinis* | 20.44 | 15.97 | PBDB |
| *Parietobalaena campiniana* | 15.0 | 13.2 | Marx and Fordyce (2015) |
| *Parietobalaena laxata* | 15.97 | 11.62 | PBDB |
| *Parietobalaena palmeri* | 16.4 | 14.5 | Marx and Fordyce (2015) |
| *Parietobalaena securis* | 15.97 | 13.82 | PBDB |
| *Parietobalaena yamaokai* | 17 | 14.9 | Marx and Fordyce (2015) |
| *Parietobalaena* sp SMNH VeF62 | 16.4 | 15.1 | Marx and Fordyce (2015) |
| Patriocetidae ChM PV2761 | 28.4 | 23.03 | PBDB |
| *Patriocetus denggi* | 28.1 | 23.03 | PBDB |
| *Patriocetus ehrlichii* | 28.1 | 23.03 | PBDB |
| *Patriocetus kazakhstanicus* | 28.1 | 23.03 | PBDB |
| *Pelocetus calvertensis* | 15.5 | 14.5 | Marx and Fordyce (2015) |
| *Pelocetus mirabilis* | 13.82 | 7.246 | PBDB |
| *Peregocetus pacificus* | 47.8 | 41.3 | Lambert et al. (2019) |
| *Peripolocetus vexillifer* | 16 | 15.2 | Marx and Fordyce (2015) |
| *Phoberodon arctirostris* | 20.44 | 15.97 | PBDB |
| *Phocaenopsis mantelli* | 19 | 15.9 | PBDB |
| *Phocaenopsis scheynensis* | 13.82 | 11.62 | PBDB |
| *Phocageneus venustus* | 20.44 | 15.97 | PBDB |
| *Phococetus vasconum* | 20.44 | 15.97 | PBDB |
| *Physeter vetus* | 15.97 | 13.82 | PBDB |
| *Physeter antiquus* | 5.33 | 2.588 | PBDB |
| Physeteridae IRSNB M 1937 | 11.608 | 5.333 | PBDB |
| *Physeterula dubusi* | 20.44 | 15.97 | PBDB |
| *Physeterula neolassicus* | 2.588 | 0.0117 | PBDB |
| *Pinocetus polonicus* | 15.97 | 13.82 | PBDB |
| *Piscobalaena nana* | 7.5 | 5.9 | Marx and Fordyce (2015) |
| *Piscocetus sacaco* | 7.246 | 5.333 | PBDB |

| | | | |
|---|---|---|---|
| *Piscolithax aenigmaticus* | 11.62 | 7.246 | PBDB |
| *Piscolithax boreios* | 7.246 | 5.333 | PBDB |
| *Piscolithax longirostris* | 7.246 | 5.333 | PBDB |
| *Piscolithax tedfordi* | 7.246 | 5.333 | PBDB |
| *Pithanodelphis cornutus* | 16.5 | 15.1 | Louwye et al. (2010) |
| *Placoziphius duboisii* | 23.03 | 15.97 | PBDB |
| *Platalearostrum hoekmani* | 3.6 | 0.8 | PBDB |
| Platanistinae MUSM 1611 | 13.2 | 12.5 | PBDB |
| Platanistoidea OU 22670 | 23.03 | 20.44 | PBDB |
| *Platylithax robusta* | 15.97 | 13.82 | PBDB |
| *Platyosphys aithai* | 41.3 | 38 | PBDB |
| *Platyosphys paulsonii* | 38 | 33.9 | PBDB |
| *Platyosphys einori* | 41.2 | 37.8 | PBDB |
| *Plesiobalaenoptera quarantellii* | 11.6 | 9.4 | Marx and Fordyce (2015) |
| *Plesiocetopsis hupschii* | 16.5 | 15.1 | Louwye et al. (2010) |
| *Plesiocetopsis notopelagicus* | 5.33 | 2.58 | PBDB |
| *Plesiocetus dyticus* | 20.44 | 15.97 | PBDB |
| *Plesiocetus giganteus* | 11.62 | 7.246 | PBDB |
| *Plesiocetus intermedius* | 11.62 | 7.246 | PBDB |
| *Plesiocetus minor* | 11.62 | 7.246 | PBDB |
| *Plesiocetus rostratus* | 11.62 | 7.246 | PBDB |
| *Plesiocetus tertius* | 11.62 | 7.246 | PBDB |
| *Plesiocetus occidentalis* | 16.3 | 13.6 | PBDB |
| *Pliokogia apenninica* | 5.08 | 5.04 | Collareta et al. (2019) |
| *Pliopontos littoralis* | 7.246 | 3.6 | PBDB |
| *Pomatodelphis bobengi* | 13.82 | 10.3 | PBDB |
| *Pomatodelphis inaequalis* | 13.82 | 4.9 | PBDB |
| *Pomatodelphis stenorhynchus* | 20.44 | 15.97 | PBDB |
| *Pomatodelphis* sp USNM 187414 | 13.82 | 11.63 | PBDB |
| *Pontistes rectifrons* | 13.82 | 11.62 | PBDB |
| *Praekogia cedrosensis* | 7.246 | 5.333 | PBDB |
| *Praemegaptera pampauensis* | 13.82 | 11.62 | PBDB |
| *Preaulophyseter gualichensis* | 23.03 | 15.97 | PBDB |
| *Prepomatodelphis korneuburgensis* | 15.97 | 13.65 | PBDB |
| *Priscophyseter typus* | 5.33 | 2.58 | PBDB |
| *Prosqualodon australis* | 23.03 | 15.97 | PBDB |
| *Prosqualodon davidis* | 23.03 | 20.44 | PBDB |
| *Prosqualodon hamiltoni* | 28.1 | 23.03 | PBDB |
| *Protocetus atavus* | 47.8 | 41.3 | PBDB |
| *Protodelphinus capellinii* | 23.03 | 20.44 | PBDB |
| *Protoglobicephala mexicana* | 3.6 | 2.588 | PBDB |
| *Protophocaena minima* | 11.62 | 7.246 | PBDB |
| *Protororqualus cuvieri* | 12.7 | 11.608 | PBDB |
| *Pseudorca yokoyamai* | 1.806 | 0.781 | PBDB |
| *Pterocetus benguelae* | 16 | 5.3 | Bianucci et al. (2007) |
| *Pterophocaena nishinoi* | 11.62 | 7.246 | PBDB |
| *Qaisracetus arifi* | 47.8 | 41.3 | PBDB |
| *Rayanistes afer* | 47.8 | 41.3 | PBDB |
| *Remingtonocetus domandaensis* | 47.8 | 41.3 | PBDB |
| *Remingtonocetus harudiensis* | 47.8 | 41.3 | PBDB |
| *Rhinostodes lovisatoi* | 11.63 | 7.246 | PBDB |
| *Rodhocetus balochistanensis* | 47.8 | 41.3 | PBDB |
| *Rodhocetus kasranii* | 46.5 | 46.5 | PBDB |
| *Rudicetus squalodontoides* | 15.97 | 7.246 | PBDB |
| *Sachalinocetus cholmicus* | 23 | 11.6 | PBDB |
| *Saghacetus osiris* | 38 | 33.9 | PBDB |
| *Salishicetus meadi* | 27.82 | 23.03 | PBDB |
| *Salumiphocaena stocktoni* | 11.62 | 7.246 | PBDB |
| *Sarmatodelphis moldavicus* | 12.7 | 11.608 | PBDB |

| | | | |
|---|---|---|---|
| *Saurocetes argentinus* | 11.62 | 7.246 | PBDB |
| *Saurocetes gigas* | 9 | 6.8 | PBDB |
| *Saurocetus gibbesii* | 33.9 | 28.1 | PBDB |
| *Scaldicetus bellunensis* | 23.03 | 20.44 | PBDB |
| *Scaldicetus caretti* | 11.62 | 1.806 | PBDB |
| *Scaldicetus degiorgii* | 15.97 | 7.246 | PBDB |
| *Scaldicetus grandis* | 11.63 | 5.33 | PBDB |
| *Scaldicetus inflatus* | 13.82 | 11.62 | PBDB |
| *Scaldicetus macgeei* | 7.246 | 5.333 | PBDB |
| *Scaldicetus minor* | 5.333 | 2.588 | PBDB |
| *Scaldicetus perpinguis* | 20.44 | 15.97 | PBDB |
| *Scaldicetus antwerpiensis* | 16.5 | 15.1 | Louwye et al. (2010) |
| *Scaldiporia vandokkumi* | 7.6 | 5 | Post et al. (2017) |
| *Scaphokogia cochlearis* | 11.62 | 7.246 | PBDB |
| Scaphokogiinae MUSM 3291 and 3405 | 11.608 | 5.332 | PBDB |
| *Schizodelphis barnesi* | 20.44 | 15.97 | PBDB |
| *Schizodelphis bogatshowi* | 13.82 | 11.62 | PBDB |
| *Schizodelphis morckhoviensis* | 15.97 | 11.62 | PBDB |
| *Schizodelphis planus* | 20.44 | 15.97 | PBDB |
| *Schizodelphis squalodontoides* | 20.44 | 15.97 | PBDB |
| *Schizodelphis sulcatus* | 20.44 | 15.97 | PBDB |
| *Schizodelphis compressus* | 11.608 | 5.332 | PBDB |
| *Semirostrum ceruttii* | 5.333 | 2.588 | PBDB |
| *Septemtriocetus bosselaersi* | 3.6 | 2.588 | PBDB |
| *Septidelphis morii* | 5.333 | 2.588 | PBDB |
| *Simocetus rayi* | 33.7 | 30.6 | PBDB |
| *Sinanodelphis izumidaensis* | 15.97 | 13.82 | PBDB |
| *Sitsqwayk cornishorum* | 28.1 | 23.03 | PBDB |
| *Sophianacetus commenticius* | 13.82 | 11.62 | PBDB |
| Squalodelphinidae USNM 475596 | 23.03 | 20.43 | PBDB |
| Squalodelphinidae UWBM 87105 | 23.03 | 20.43 | PBDB |
| *Squalodelphis fabianii* | 23.03 | 20.44 | PBDB |
| *Squalodon antverpiensis* | 16.5 | 15.1 | Louwye et al. (2010) |
| *Squalodon atlanticus* | 15.97 | 13.82 | PBDB |
| *Squalodon barbarus* | 28.1 | 23.03 | PBDB |
| *Squalodon bariensis* | 23.03 | 15.97 | PBDB |
| *Squalodon bellunensis* | 23.04 | 20.44 | PBDB |
| *Squalodon bordae* | 20.44 | 15.97 | PBDB |
| *Squalodon calvertensis* | 23.03 | 20.44 | PBDB |
| *Squalodon catulli* | 23.03 | 20.44 | PBDB |
| *Squalodon dalpiazi* | 20.44 | 15.97 | PBDB |
| *Squalodon grateloupii* | 28.1 | 23.03 | PBDB |
| *Squalodon hypsispondylus* | 28.1 | 23.03 | PBDB |
| *Squalodon imperator* | 13.82 | 11.62 | PBDB |
| *Squalodon linzianus* | 28.1 | 23.03 | PBDB |
| *Squalodon melitensis* | 20.44000 | 15.97 | PBDB |
| *Squalodon meyeri* | 20.44 | 15.97 | PBDB |
| *Squalodon molassicus* | 20.44 | 15.97 | PBDB |
| *Squalodon peregrinus* | 23.03 | 20.44 | PBDB |
| *Squalodon protervus* | 15.97 | 11.62 | PBDB |
| *Squalodon servatus* | 20.44 | 15.97 | PBDB |
| *Squalodon tiedemani* | 23.03 | 20.43 | PBDB |
| *Squalodon vocontiorum* | 20.44 | 15.97 | PBDB |
| *Squalodon whitmorei* | 20.44 | 15.97 | PBDB |
| *Squalodon mirabilis* | 15.97 | 11.608 | PBDB |
| *Squalodon* sp MAUL 8 | 20.43 | 15.97 | PBDB |
| *Squalodon* sp OU 21798 | 27.82 | 23.03 | PBDB |
| Squalodontidae ChM PV4961 | 27.82 | 23.03 | PBDB |
| Squalodontidae ChM PV4991 | 27.82 | 23.03 | PBDB |

| | | | |
|---|---|---|---|
| Squalodontidae OU 21798 | 27.82 | 23.03 | Fordyce (1994) |
| *Squaloziphius emlongi* | 23.03 | 20.44 | PBDB |
| *Stenasodelphis russellae* | 11.62 | 7.246 | PBDB |
| *Stenella rayi* | 5.333 | 3.6 | PBDB |
| *Stromerius nidensis* | 38 | 33.9 | PBDB |
| *Sulaimanitherium dhanotri* | 41.3 | 38 | PBDB |
| *Sulakocetus dagestanicus* | 28.1 | 23.03 | PBDB |
| *Supayacetus muizoni* | 41.3 | 38 | PBDB |
| *Tagicetus joneti* | 13.82 | 11.62 | PBDB |
| *Taikicetus inouei* | 15.97 | 11.62 | PBDB |
| *Takracetus simus* | 47.8 | 41.3 | PBDB |
| *Tangaroasaurus kakanuiensis* | 15.97 | 13.82 | PBDB |
| *Thalassocetus antwerpiensis* | 16.5 | 15.1 | Louwye et al. (2010) |
| Thalassotherii USNM 187416 | 15.97 | 13.63 | PBDB |
| *Thinocetus arthritus* | 13.5 | 13.4 | Marx and Fordyce (2015) |
| *Tiphyocetus temblorensis* | 16 | 15.2 | Marx and Fordyce (2015) |
| *Titanocetus sammarinensis* | 16.4 | 14.7 | Marx and Fordyce (2015) |
| *Tiucetus rosae* | 11 | 9 | Marx et al. (2017) |
| *Tlaxcallicetus guaycurae* | 28.4 | 23.03 | PBDB |
| *Tlaxcallicetus* sp MU EcSj5/18/95 | 28.4 | 23.03 | PBDB |
| *Togocetus traversei* | 47.8 | 41.3 | PBDB |
| *Tohoraata raekohao* | 27.3 | 25.2 | PBDB |
| *Tohoraata waitakiensis* | 27.3 | 25.2 | PBDB |
| *Toipahautea waitaki* | 27.3 | 25.2 | PBDB |
| *Tokarahia kauaeroa* | 27.3 | 25.2 | PBDB |
| *Tokarahia lophocephalus* | 27.3 | 25.2 | PBDB |
| *Tranatocetus argillarius* | 11.62 | 7.246 | PBDB |
| *Tranatocetus maregermanicum* | 8.1 | 7.5 | Marx et al. (2019) |
| *Tupelocetus palmeri* | 41.3 | 38 | Gibson et al. (2018) |
| *Tursiops brochii* | 3.6 | 2.588 | PBDB |
| *Tursiops capellinii* | 5.333 | 2.588 | PBDB |
| *Tursiops osennae* | 3.6 | 2.588 | PBDB |
| *Tursiops astensis* | 3.5 | 3 | PBDB |
| *Tusciziphius atlanticus* | 6.1 | 4.4 | PBDB |
| *Tusciziphius crispus* | 4.12 | 3.84 | PBDB |
| *Uranocetus gramensis* | 8.6 | 7.5 | Marx and Fordyce (2015) |
| *Urkudelphis chawpipacha* | 26 | 24 | Tanaka et al. (2017) |
| *Vampalus sayasanicus* | 15.97 | 5.33 | PBDB |
| *Vanbreenia trigonia* | 15.97 | 13.82 | PBDB |
| *Waharoa ruwhenua* | 25.4 | 25.3 | Marx and Fordyce (2015) |
| *Waipatia maerewhenua* | 27.2 | 24.5 | Marx and Fordyce (2015) |
| *Waipatia hectori* | 27.3 | 25.2 | PBDB |
| *Whakakai waipata* | 27.3 | 25.2 | PBDB |
| *Wimahl chinookensis* | 23.03 | 15.97 | PBDB |
| Xenorophidae ChM PV2758 | 27.82 | 23.03 | PBDB |
| Xenorophidae ChM PV4746 | 27.82 | 23.03 | PBDB |
| Xenorophidae ChM PV5711 | 27.82 | 23.03 | PBDB |
| *Xenorophus sloanii* | 33.9 | 28.1 | PBDB |
| *Xenorophus* sp ChM PV4823 | 30 | 27 | PBDB |
| *Xhosacetus hendeysi* | 16 | 5.3 | Bianucci et al. (2007) |
| *Xiphiacetus bossi* | 20.44 | 15.97 | PBDB |
| *Xiphiacetus cristatus* | 16.5 | 15.1 | Louwye et al. (2010) |
| *Yamatocetus canaliculatus* | 29.2 | 28.1 | Marx and Fordyce (2015) |
| *Yaquinacetus meadi* | 24 | 19.2 | Lambert et al. (2018b) |
| *Zarhachis crassangulum* | 15.97 | 11.62 | PBDB |
| *Zarhachis flagellator* | 23.03 | 20.44 | PBDB |
| *Zarhachis tysonii* | 11.62 | 7.246 | PBDB |
| *Zarhachis velox* | 15.97 | 13.82 | PBDB |
| *Zarhinocetus donnamatsonae* | 20.44 | 13.82 | PBDB |

| | | | |
|---|---|---|---|
| *Zarhinocetus errabundus* | 15.97 | 11.62 | PBDB |
| Ziphiidae KNM LP52956 | 23.3 | 16.1 | PBDB |
| *Ziphiodelphis abeli* | 23.03 | 20.44 | PBDB |
| *Ziphiodelphis sigmoideus* | 23.03 | 20.44 | PBDB |
| *Ziphirostrum marginatum* | 9.5 | 7.5 | PBDB |
| *Ziphirostrum recurvus* | 15.97 | 5.33 | PBDB |
| *Ziphirostrum turninense* | 11.6 | 2.588 | PBDB |
| *Ziphius compressus* | 2.588 | 1.806 | PBDB |
| *Zygiocetus nartorum* | 7.246 | 5.333 | PBDB |
| *Zygophyseter varolai* | 10.5 | 8.14 | Bianucci and Landini (2006) |
| *Zygorhiza kochii* | 38 | 33.9 | PBDB |
